# Discovery of ILT3 (LILRB4) Small Molecule Inhibitors by Affinity Selection-Mass Spectrometry Reveals Druggability of a Neuroimmune Checkpoint in Alzheimer’s Disease

**DOI:** 10.64898/2026.06.05.730337

**Authors:** Somaya A. Abdel-Rahman, Natarajan Arul Murugan, Moustafa T. Gabr

## Abstract

Leukocyte immunoglobulin-like receptor B4 (LILRB4/ILT3) is an emerging neuroimmune checkpoint that restricts microglial activation and amyloid clearance in Alzheimer’s disease (AD) through ApoE-dependent signaling. Here, we establish ILT3 as a tractable small molecule target using affinity selection-mass spectrometry (AS-MS) to identify direct binders. Biophysical validation confirmed high-affinity engagement, with **LT12** exhibiting nanomolar binding by MST and SPR. Computational modeling and mutagenesis defined a discrete ILT3 binding pocket, revealing a distributed interaction network critical for ligand engagement. Targeting ILT3 disrupted the ILT3-ApoE interaction, with **LT12** showing potent inhibition in orthogonal biochemical assays. In human iPSC-derived microglia, ILT3 modulation attenuated SHP1/2 signaling, suppressed NF-κB activation, reduced IL-1β secretion, and restored Aβ uptake. In vivo, pharmacological targeting of ILT3 improved cognition, reduced amyloid burden, and attenuated neuroinflammation in 5xFAD mice. Together, these findings validate ILT3 as a druggable neuroimmune checkpoint and support its therapeutic targeting in AD.

## INTRODUCTION

Alzheimer’s disease (AD) continues to represent a profound unmet medical need, as currently available disease-modifying treatments are largely confined to amyloid-directed antibodies that offer limited clinical benefit and are hindered by suboptimal brain exposure and safety liabilities.^1-5^ A growing body of evidence highlights microglia as key regulators of amyloid-β (Aβ) pathology, with disease progression increasingly understood as a consequence of dysregulated signaling networks that govern microglial activation states.^6-10^ These cells operate under a finely tuned balance of stimulatory and inhibitory inputs, mediated by receptors such as triggering receptor expressed on myeloid cells 2 (TREM2) and suppressive pathways involving CD33 and members of the leukocyte immunoglobulin-like receptor (LILR) family, which together influence phagocytosis and inflammatory responses.^4,5^ Recent work, including our own, has demonstrated that small-molecule modulation of TREM2 can reprogram microglial activity, underscoring the therapeutic potential of targeting neuroimmune checkpoint pathways.^11-14^

Among inhibitory regulators, leukocyte immunoglobulin-like receptor B4 (LILRB4, also known as ILT3) has recently gained attention as a critical microglial checkpoint that restricts Aβ clearance through ApoE-dependent mechanisms.^15-22^ ILT3 (LILRB4) expression is markedly elevated in microglia from AD patients, particularly within plaque-associated subsets, and shows strong correlation with ApoE levels and pathological features such as phosphorylated tau.^23^ Upon activation, ILT3 signals through immunoreceptor tyrosine-based inhibitory motifs (ITIMs), leading to recruitment of SHP1 and SHP2 phosphatases and suppression of cytoskeletal dynamics and phagocytic machinery required for efficient plaque removal.^16,19^ In preclinical models, antibody-mediated inhibition of ILT3 enhances microglial activation, promotes Aβ uptake, and reduces amyloid burden by approximately 50%, alongside attenuation of interferon-related programs and remodeling of plaque-associated microglial populations.^23^ Collectively, these findings position ILT3 as a promising therapeutic target in AD and highlight the ILT3-ApoE interaction as a defined and druggable interface for small molecule intervention. However, targeting this interaction remains challenging due to its non-enzymatic, protein-protein interface and the lack of robust activity-based screening assays, particularly in the context of identifying brain-permeable small molecules suitable for central nervous system (CNS) applications.

To address this challenge, we applied affinity selection mass spectrometry (AS-MS), a label-free, solution-phase platform capable of identifying protein–ligand interactions with high specificity and throughput.^24,25^ In contrast to conventional high-throughput screening (HTS) approaches that depend on enzymatic activity or fluorescence-based readouts, AS-MS enables the direct detection of ligand binding independent of functional output.^26^ Adapting this approach to ILT3, a non-enzymatic and structurally flexible immune receptor, required methodological optimization. We developed a size exclusion chromatography (SEC)-coupled AS-MS workflow, systematically refining protein preparation, buffer composition, library diversity, and chromatographic resolution to enable selective enrichment of ILT3-ligand complexes followed by high-resolution mass spectrometric identification. Compounds emerging from the primary screen of a CNS-focused chemical library were progressed through a multi-stage validation cascade.

Here, we report the discovery of small molecule ILT3 binders identified through AS-MS screening that directly engage the receptor and modulate its immunoregulatory function. Biophysical validation and functional studies in human microglia demonstrate that these compounds enhance phagocytic activity and attenuate inhibitory signaling pathways associated with ILT3 activation. A multi-tiered validation framework, incorporating orthogonal biophysical methods and complementary cellular assays, confirmed target engagement and functional modulation across systems. Pharmacokinetic (PK) and in vivo evaluations demonstrate brain penetration and are associated with improved cognitive performance, reduced amyloid burden, and attenuation of neuroinflammation in the 5xFAD mouse model. Together, these findings establish ILT3 as a tractable target for small molecule modulation and provide a foundation for further optimization and therapeutic development.

## RESULTS AND DISCUSSION

### AS-MS Screening to Identify ILT3-targeted compounds

To identify small molecule ligands capable of engaging ILT3, we implemented a two-dimensional AS-MS platform incorporating SEC to enable detection of protein-ligand interactions under native solution conditions (Figure 1A). In this workflow, recombinant ILT3 extracellular domain was incubated with pooled small molecule libraries, followed by SEC separation to isolate protein-associated fractions. These fractions were subsequently denatured and analyzed by high-resolution LC-MS to identify ligands retained through binding interactions.

**Figure 1.**
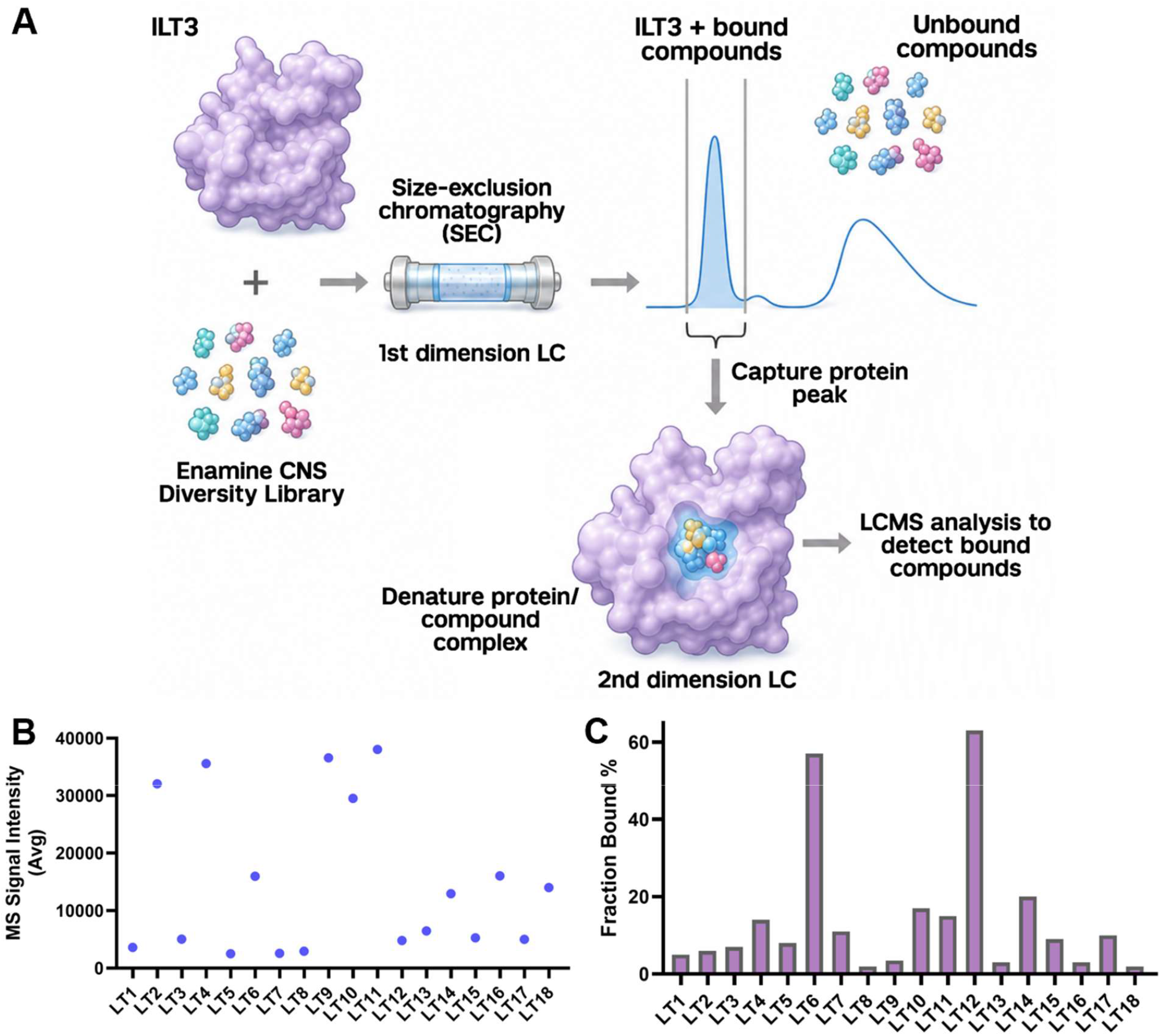
Two-dimensional SEC-coupled AS-MS screening enables identification of ILT3-binding small molecules. **(A)** Overview of the SEC-coupled affinity selection–mass spectrometry workflow used to detect ligands that associate with ILT3 under solution-phase conditions. A library of 5,441 compounds was screened in pooled format in the presence of ILT3, followed by SEC separation and LC-MS analysis to identify retained ligands. **(B)** Broad distribution of MS signal intensities of the selected 18 hits. Candidate selection was performed using stringent filtering criteria, including mass accuracy within 5 ppm, absence of signal in compound-only controls (P-), enrichment defined by a P+/P− intensity ratio ≥3, consistent retention time, well-resolved isotopic distribution, and signal intensity exceeding 1,000 counts. **(C)** Fraction bound (Fb%) values for compounds validated in the secondary screen using 2.5 µM ILT3. Compounds **LT6** and **LT12** exhibited the highest enrichment (Fb% >50%), consistent with strong binding to ILT3.

In the primary screening campaign, approximately 5,441 compounds from the Enamine CNS Diversity Library were evaluated at a final concentration of 1 µM in the presence of 1 µM ILT3. Compounds were screened in pooled format, with each pool containing ~1,100 molecules. SEC/UV chromatograms revealed well-defined protein peaks in LILRB4-containing samples (P1+ and P2+), which were absent in compound-only control samples (P-), confirming effective separation of protein-ligand complexes from unbound compounds. LC-MS analysis identified 36 preliminary binders based on stringent selection criteria, including mass accuracy within 5 ppm, absence of signal in the P-control, enrichment defined by a P+/P-intensity ratio ≥3, consistent retention time, well-resolved isotopic patterns, and signal intensity exceeding 1,000 counts.

To assess binding specificity and reproducibility, these 36 candidates were subjected to a secondary screening round using an increased protein concentration (2.5 µM ILT3) and reduced pool complexity (~8 compounds per pool). This follow-up screen yielded 18 reproducible hits, corresponding to a hit rate of 0.33%, all of which satisfied the same selection thresholds. The chemical structures of these 18 compounds are provided in Table S1. As shown in Figure 1B, the validated hits displayed a broad distribution of MS signal intensities, with several compounds exhibiting particularly strong enrichment. Binding was further quantified using fraction bound (Fb%), representing the proportion of each ligand associated with the protein-containing fraction. As illustrated in Figure 1C, compounds **LT6** and **LT12** emerged as the most prominent binders, with Fb% values of 57%, and 63%, respectively. Collectively, these data demonstrate the robustness and selectivity of the AS-MS platform for identifying ILT3-binding small molecules and provide a prioritized set of chemically tractable candidates for downstream biophysical validation, functional characterization, and structure-guided optimization targeting ILT3-mediated neuroimmune signaling.

#### Validation of Hit Compounds

To validate binding of compounds identified from the SEC-coupled AS-MS screen, all 18 confirmed hits were evaluated using a fluorescence-based assay on the Dianthus platform, which detects molecular interactions through temperature-induced changes in fluorescence intensity, referred to as temperature-related intensity change (TRIC) (Figure 2A). Compounds were tested at a single concentration of 50 µM in PBST buffer (PBS pH 7.4 containing 0.05% Tween-20) supplemented with 1% DMSO. ILT3 binding signals were quantified as normalized fluorescence (F_norm_) relative to DMSO-treated controls. Compounds producing a change in fluorescence (ΔF_norm_) of at least 5 units and showing reproducible responses across replicates (n = 5) were designated as preliminary hits (Figure 2A). Using these criteria, 5 compounds (Figure 2B) were identified in the primary screen as potential ILT3 hits.

**Figure 2.**
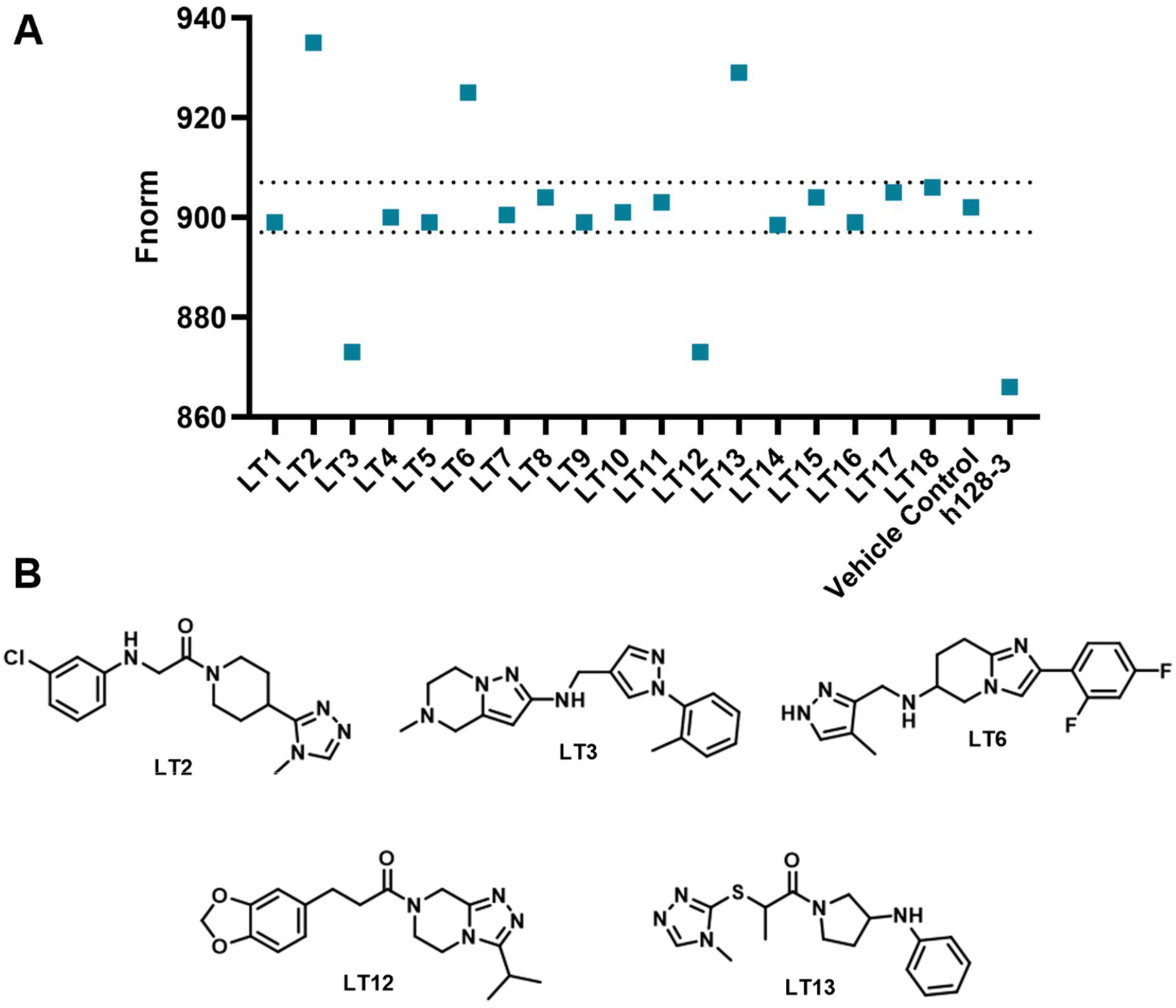
**(A)** Single-concentration (at 50 μM) TRIC-based screening of 18 compounds from AS-MS screening against the ILT3 extracellular domain. Binding responses are reported as normalized fluorescence (F_norm_). The dashed line denotes the hit cutoff (ΔF_norm_ ≥ 5), and compounds exceeding this threshold were designated as preliminary binders. **(B)** Chemical structures of 5 preliminary hits from AS-MS screening as potential ILT3 binders.

To quantify binding affinity, these compounds were further evaluated using microscale thermophoresis (MST). While both Dianthus and MST rely on TRIC-based detection, MST provides enhanced sensitivity, improved control of thermal gradients, and reduced sample consumption, making it well suited for quantitative analysis. Thermophoresis signals were fitted to generate binding curves and determine equilibrium dissociation constants (Kd). Two compounds exhibited clear concentration-dependent binding behavior. The measured Kd values were: **LT6** (2.92 ± 0.84 µM, Figure S1) and **LT12** (41.3 ± 6.8 nM, Figure 3A). To further validate binding using an orthogonal biophysical approach, the most potent compound, **LT12**, was evaluated by surface plasmon resonance (SPR). SPR analysis revealed a concentration-dependent binding response consistent with a direct interaction between **LT12** and ILT3, yielding a Kd of 81.3 ± 9.4 nM (Figure 3B). The close agreement between MST and SPR measurements supports the robustness of the observed interaction and confirms **LT12** as a high-affinity binder of ILT3.

**Figure 3.**
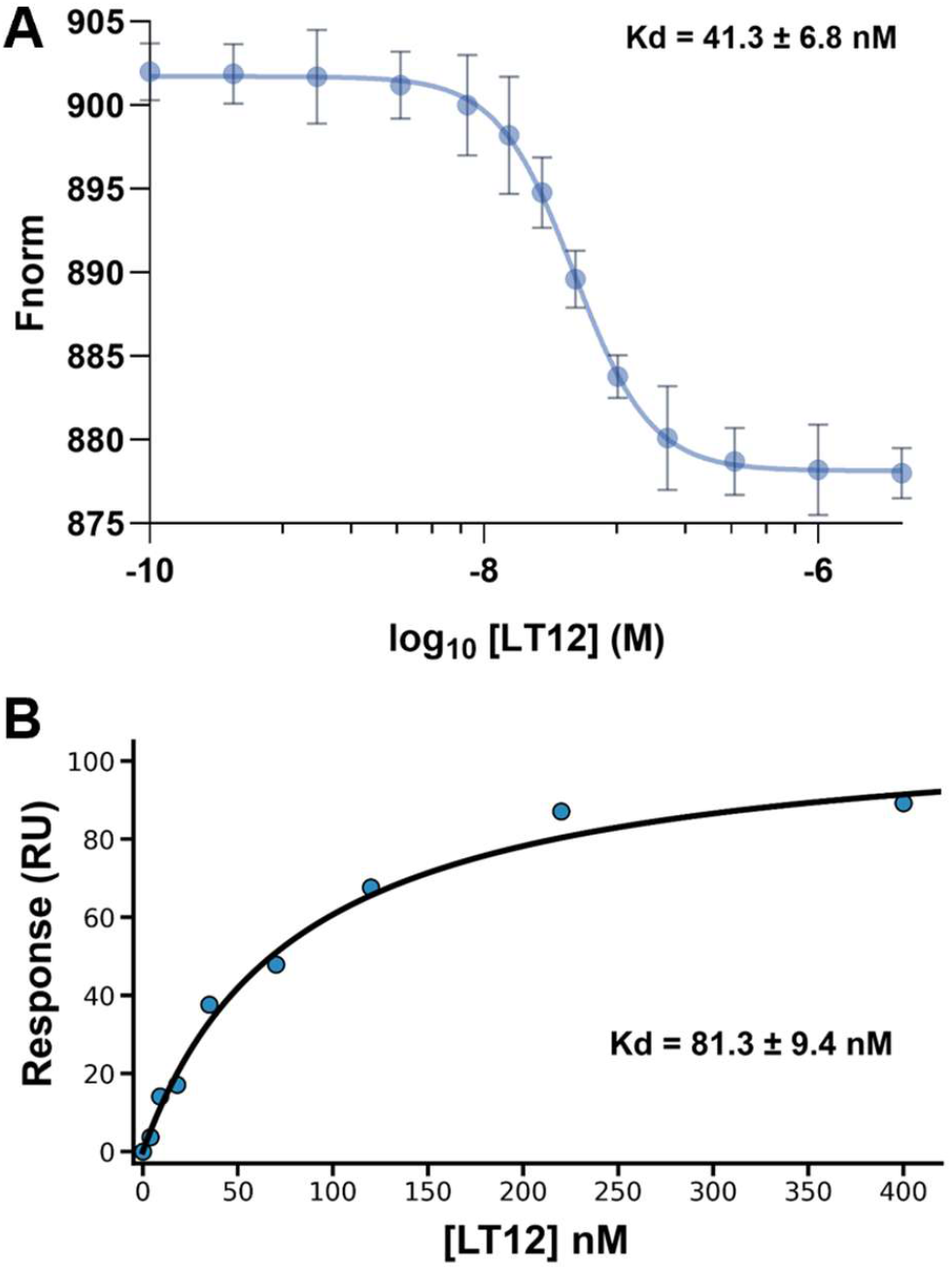
MST and SPR validation of LT12 binding to ILT3. **(A)** MST dose-response curve for **LT12**, indicating strong and saturable binding to ILT3 (Kd = 41.3 ± 6.8 nM). Data are shown as mean ± SD (n = 5). **(B)** Orthogonal validation of compound **LT12** binding by SPR, demonstrating a concentration-dependent response consistent with a hyperbolic binding profile (Kd = 81.3 ± 9.4 nM, n = 3).

### Computational Analysis and Site-Directed Mutagenesis Validation of Ligand Binding

Blind docking identified energetically favorable binding poses for both ligands within ILT3 (PDB ID: 3P2T), with **LT6** and **LT12** occupying distinct regions of the receptor (Figure S2 and Figure 4A, respectively). For **LT6**, the predicted binding mode shows a protonated amine forming a hydrogen bond with Pro89, with the ligand positioned in a pocket defined by Leu90, Glu91, Pro89, and Val93, while the difluorophenyl group extends into a hydrophobic region (Figure S3). In contrast, the predicted pose of **LT12** features a hydrogen bond between a carbonyl group and Asn20, along with hydrophobic contacts involving Val15, Ile16, and Pro12, resulting in a broader interaction footprint within the binding pocket (Figure 4B). Despite similar docking scores (-8.03 and -8.12 kcal/mol), these computationally predicted interactions indicate a localized interaction pattern for **LT6** and a broader residue engagement for **LT12**.

**Figure 4.**
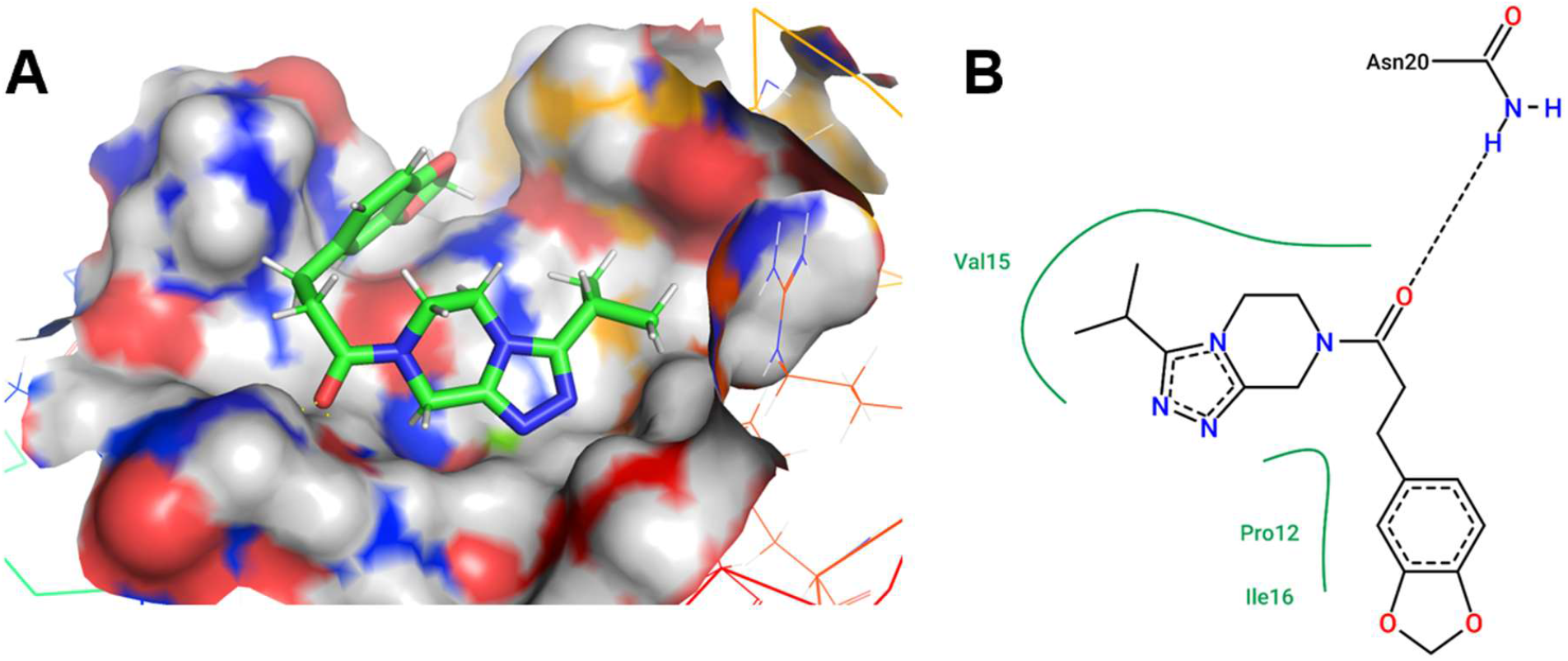
Computationally predicted binding mode of LT12 in ILT3. **(A)** Docked pose of **LT12** (green) within the ILT3 binding pocket, shown on the protein surface colored by physicochemical properties. **(B)** Two-dimensional interaction map highlighting key predicted interactions between **LT12** and ILT3. Dashed lines represent hydrogen bonds and solid green lines represent hydrophobic contacts.

Molecular dynamics (MD) simulations (Figure 5A) show that both **LT6** and **LT12** remain bound to ILT3 over the 80 ns trajectory. **LT6** exhibits RMSD values primarily in the range of ~1-3 Å, whereas **LT12** shows higher RMSD values of ~2-4 Å. Residue-wise free energy decomposition (Figure 5B) reveals distinct interaction profiles for the two ligands, with **LT6** displaying limited contributions from a small number of residues, while **LT12** engages a broader set of residues with more distributed negative ΔG contributions. Consistent with these observations, MM/GBSA analysis (Table S2) indicates that **LT12** has a more favorable total binding free energy (ΔG_total_ = -25.1 kcal/mol) compared to **LT6** (-11.4 kcal/mol). This difference is primarily associated with a stronger van der Waals contribution for **LT12**, whereas the large electrostatic term observed for **LT6** is offset by an unfavorable polar solvation contribution.

**Figure 5.**
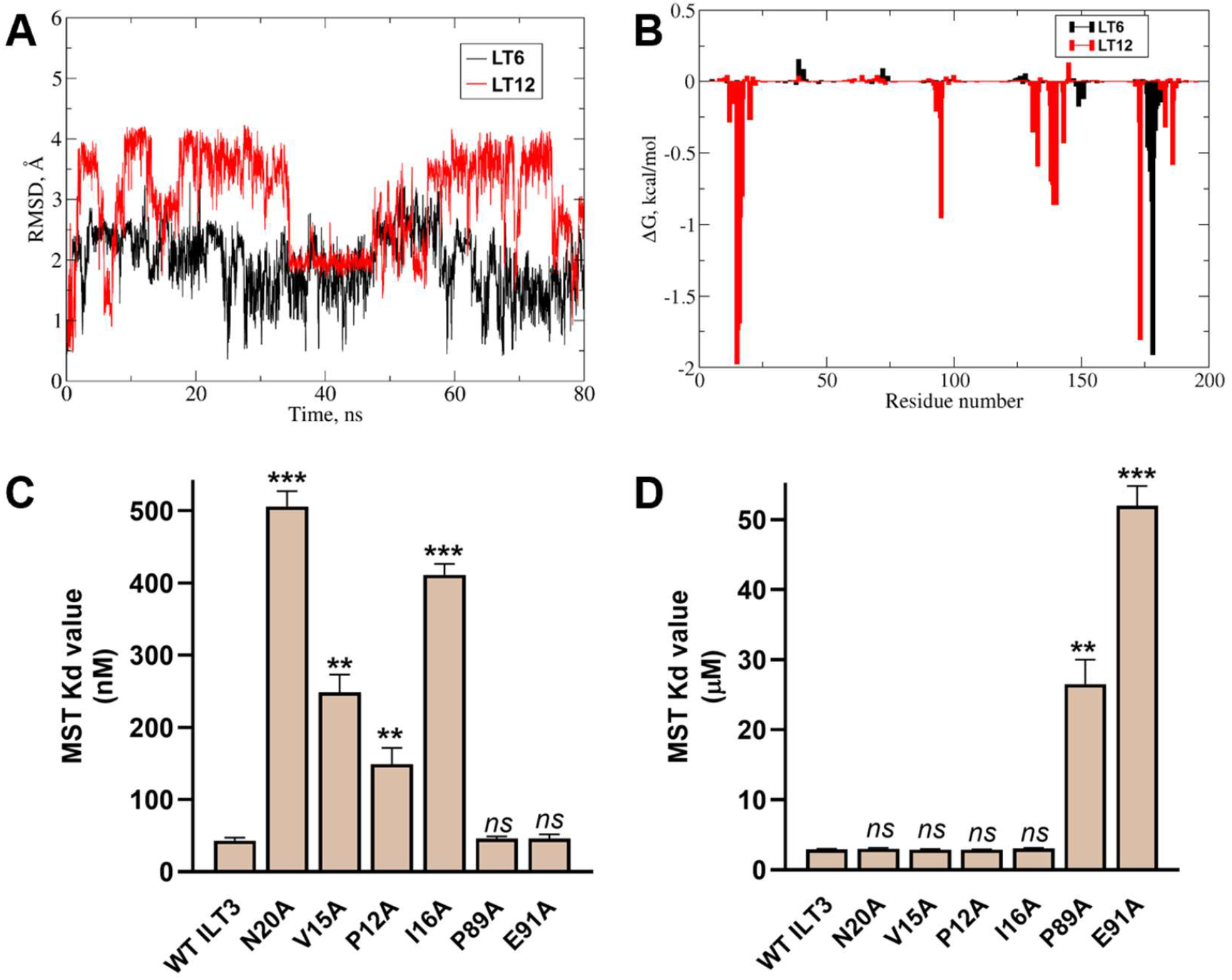
Computational and experimental validation of distinct binding modes for LT6 and LT12. **(A)** Root-mean-square deviation (RMSD) of **LT6** (black) and **LT12** (red) bound to ILT3 over 80 ns MD simulations. **(B)** Residue-wise decomposition of binding free energies (ΔG, kcal/mol per residue) from MM/GBSA analysis for **LT6** (black) and **LT12** (red). **(C)** MST-derived Kd values (nM) for **LT12** binding to wild-type (WT) ILT3 and indicated mutants. **(D)** MST-derived Kd values (µM) for **LT6** binding to WT ILT3 and indicated mutants. In contrast to **LT12, LT6** binding is selectively affected by P89A and E91A, with no significant effects observed for mutations in the **LT12**-associated residues. Data are presented as mean ± SD (n=5). Statistical significance is indicated as *ns*, not significant; **, *p* < 0.01; ***, *p* < 0.001 (vs WT ILT3).

To experimentally validate the binding mode of **LT12**, site-directed mutagenesis was performed on residues identified in the predicted **LT12** interaction region, including Asn20Ala (N20A), Val15Ala (V15A), Ile16Ala (I16A), and Pro12Ala (P12A), along with residues from the **LT6**-associated region (Pro89Ala (P89A) and Glu91Ala (E91A)). Relative to wild-type (WT) ILT3, mutations in the **LT12**-associated residues resulted in significant increases in apparent MST Kd values, with N20A and I16A producing the largest effects, followed by V15A and P12A (Figure 5C). In contrast, mutations in the **LT6**-associated residues (P89A and E91A) showed no significant effect on **LT12** binding (Figure 5D). These results support the computationally predicted **LT12** binding site and indicate that interactions within the Asn20/Val15/Ile16/Pro12 region are critical for ligand affinity.

### Inhibition of the ILT3-ApoE Interaction by LT6 and LT12

Following confirmation that **LT6** and **LT12** bind directly to ILT3, we next examined whether this binding translates into functional interference with ligand engagement. ApoE was selected as the ligand given its reported interaction with ILT3 and its relevance to immune modulation in AD.^23^ To address this, the inhibitory activity of both compounds was evaluated using complementary biochemical assays. In an ELISA-based format, the ILT3 extracellular domain was immobilized and incubated with its ligand in the presence of increasing concentrations of **LT6** or **LT12. LT12** exhibited potent inhibition, with an IC_50_ of 107 ± 8.5 nM (Figure 6A), whereas **LT6** showed more modest activity, with an IC_50_ of 15.6 ± 1.8 µM (Figure S4), consistent with their respective binding affinities determined by MST.

**Figure 6.**
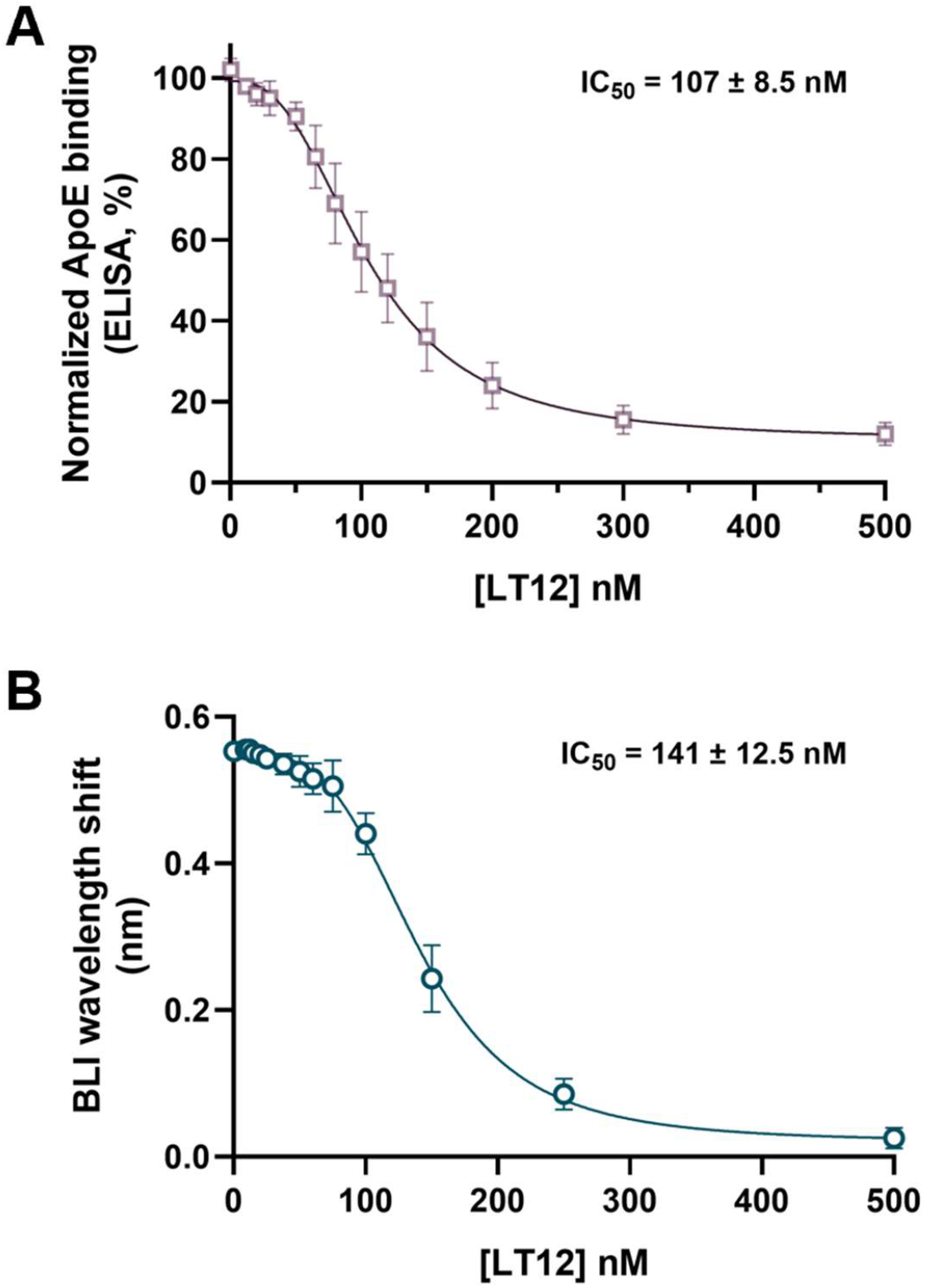
Inhibition of the ILT3–ApoE interaction by LT6 and LT12. **(A)** Dose-response curves from ELISA showing inhibition of ApoE binding to immobilized ILT3 in the presence of increasing concentrations of **LT12. (B)** BLI analysis of ApoE binding to ILT3 in the presence of **LT12**. Data are presented as mean ± SD (n=5).

To confirm these findings using an independent method, competitive binding experiments were performed by biolayer interferometry (BLI), enabling real-time monitoring of ligand association with ILT3. Consistent with the ELISA results, **LT12** reduced the binding response in a concentration-dependent manner with an IC_50_ of 141 ± 12.5 nM (Figure 6B), while **LT6** displayed weaker inhibition with an IC_50_ of 33.5 ± 4.2 µM (Figure S5). The agreement between ELISA and BLI across independent assay formats supports a direct effect of both compounds on receptor-ligand interaction.

### Pharmacological Disruption of ILT3-ApoE Signaling Restores Microglial Function

To evaluate whether inhibition of the ILT3-ApoE interaction translates into functional effects in disease-relevant systems, **LT12** (the most potent ILT3 inhibitor from this work) was assessed in human iPSC-derived microglia under Aβ-driven conditions. Cells were treated with Aβ_42_ oligomers in the presence or absence of ApoE to model ligand-dependent ILT3 activation, followed by exposure to **LT12** (0.5, 2, and 5 µM).

We first examined proximal receptor signaling by measuring phosphorylation of SHP1 and SHP2, phosphatases recruited to ITIM-containing receptors such as ILT3. Co-treatment with Aβ_42_ and ApoE resulted in a pronounced increase in phospho-SHP1 and phospho-SHP2 compared to vehicle and Aβ_42_ alone (Figures 7A and S6, respectively), consistent with activation of ILT3-mediated inhibitory signaling. **LT12** treatment reduced SHP1 and SHP2 phosphorylation in a concentration-dependent manner, with partial attenuation at 0.5 µM and more substantial suppression at 2 and 5 µM (Figures 7A and S6), indicating effective disruption of ligand-induced receptor signaling in microglia. We next evaluated downstream inflammatory signaling by quantifying NF-κB activation. Stimulation with Aβ_42_ increased NF-κB activity, which was further enhanced by ApoE co-treatment (Figure 7B). **LT12** treatment decreased NF-κB activation in a dose-dependent fashion, with higher concentrations restoring activity toward baseline levels (Figure 7B). The parallel reduction in SHP1/2 phosphorylation and NF-κB signaling supports a mechanistic link between ILT3 engagement and downstream inflammatory pathways.

**Figure 7.**
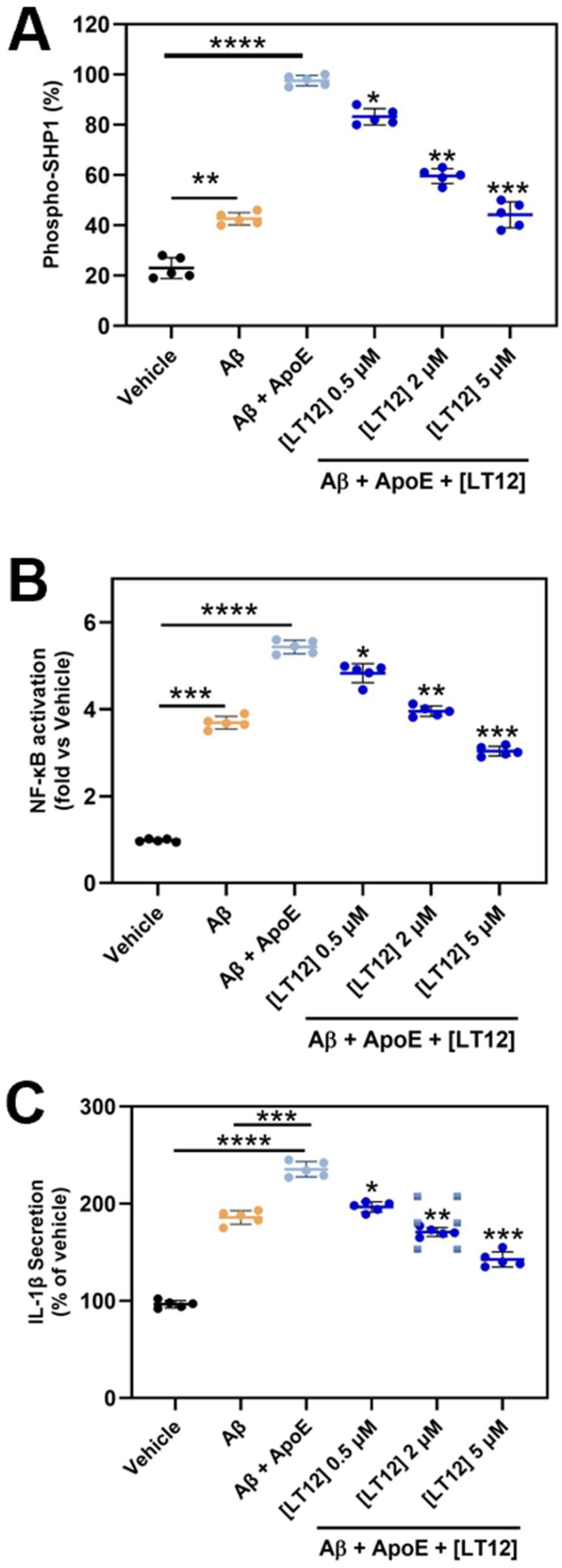
LT12 inhibits ILT3-mediated signaling in iPSC-derived human microglia. **(A)** Phosphorylation of SHP1 following stimulation with Aβ_42_ and ApoE. ApoE co-treatment enhanced SHP1 phosphorylation relative to Aβ_42_ alone, consistent with activation of ILT3-dependent signaling. **LT12** reduced phospho-SHP1 levels in a concentration-dependent manner. **(B)** NF-κB activation (fold-change vs vehicle). Aβ_42_ increased NF-κB activity, which was further elevated by ApoE. LT12 treatment attenuated NF-κB activation in a dose-dependent manner. **(C)** IL-1β secretion (% of vehicle). Aβ_42_ induced IL-1β release, which was further enhanced in the presence of ApoE. **LT12** significantly reduced IL-1β secretion across the tested concentrations. Data are presented as mean ± SD (n = 5). Statistical significance was determined by one-way ANOVA with appropriate multiple-comparisons tests. Comparisons between Aβ_42_ and vehicle, and between Aβ_42_ + ApoE and control conditions, are indicated where appropriate. For compound-treated groups, statistical comparisons were performed relative to the Aβ_42_ + ApoE condition unless otherwise stated. \**p* < 0.05, \*\**p* < 0.01, \*\*\**p* < 0.001, and \*\*\*\**p* < 0.0001.

To determine functional consequences, we measured secretion of the pro-inflammatory cytokine IL-1β. Exposure to Aβ_42_ increased IL-1β release, which was further amplified in the presence of ApoE (Figure 7C). **LT12** significantly reduced IL-1β secretion across all tested concentrations, with the strongest effects observed at 2 and 5 µM, consistent with its impact on upstream signaling events. Given the central role of microglia in amyloid clearance, Aβ_42_ uptake was evaluated as a functional readout. While Aβ_42_ alone did not significantly alter uptake, ApoE co-treatment markedly reduced Aβ internalization (Figure S7), indicative of impaired microglial function. **LT12** treatment restored Aβ uptake in a concentration-dependent manner, with significant recovery at 2 and 5 µM (Figure S7).

Importantly, no significant changes in cell viability were observed across all conditions (Figure S8), indicating that the observed effects are not attributable to cytotoxicity. Together, these findings demonstrate that pharmacological disruption of ILT3-ApoE signaling by **LT12** restores microglial signaling and functional responses under Aβ-driven conditions. When considered alongside biochemical and biophysical characterization, these results establish a direct link between target engagement and functional rescue in human microglial models, supporting small molecule-based inhibition of ILT3 as a viable pathway for modulating neuroimmune dysfunction in AD.

### PK Profiling of LT12

To characterize the developability of **LT12**, we performed a comprehensive in vitro ADME and safety assessment (Table 1). **LT12** exhibited a balanced physicochemical profile, with a LogD7.4 value of 2.24, consistent with moderate lipophilicity and supportive of efficient membrane partitioning without compromising aqueous compatibility. This profile was reflected in its solubility behavior, as **LT12** demonstrated favorable kinetic solubility in PBS containing 1% DMSO (130.5 µM) and maintained high solubility in biorelevant FaSSIF media (101.5 µM), indicating favorable compatibility with gastrointestinal-like conditions.

**Table 1.** In vitro PK profile of LT12.

| PK parameter | LT12 |
| --- | --- |
| LogD<sub>7.4</sub><sup>a</sup> | 2.24 |
| Kinetic Solubility<br>1% DMSO/PBS<br>[μM]<sup>b</sup> | 130.5 |
| Solubility in FaSSIF<br>[μM] | 101.5 |
| PAMPA $P_{app}$ [cm/s] | $4.6 \times 10^{-6}$ |
| Stability in<br>simulated gastric<br>fluid (pH 1.2, 2h) | >92% |
| Stability in<br>simulated<br>intestinal fluid (pH<br>6.8, 2h) | >90% |
| t<sub>1/2</sub> [min] mouse liver microsomes<sup>c</sup> / CL<sub>int</sub><sup>d</sup> | 118.4 ± 6.7 / 15.8 ± 1.3 |
| t<sub>1/2</sub> [min] human liver microsomes<sup>c</sup> / CL<sub>int</sub><sup>d</sup> | 132.6 ± 7.9 / 12.7 ± 1.1 |
| t<sub>1/2</sub> [min] mouse plasma | >180 |
| t<sub>1/2</sub> [min] human plasma | >180 |
| t<sub>1/2</sub> [min] PBS | >300 |
| PPB human [%]<sup>c</sup> | 94.2 ± 0.61 |
| IC<sub>50</sub> against WI-38 [μM] | >100 |
| IC<sub>50</sub> against HS 27 [μM] | >100 |
| IC<sub>50</sub> against hERG [μM] | >100 |
| CYP3A4 inhibition (% at 10 μM) | 5.9 |
| CYP2D6 inhibition (% at 10 μM) | 2.8 |
| CYP2C9 inhibition (% at 10 μM) | 4.7 |
| CYP2C19 inhibition (% at 10 μM) | 3.5 |

<sup>a</sup>LogD (pH = 7.4) the distribution coefficient at physiological pH 7.4. <sup>b</sup>Kinetic solubility measured in 1% DMSO/PBS (pH 7.4) by UV-vis spectrophotometry (n=3). <sup>c</sup>Values are mean ± SD (n=3). <sup>d</sup>Intrinsic clearance expressed in μL/(min × mg) protein. Values are mean ± SD (n=3).

Membrane permeability, evaluated using PAMPA, showed a *P*_*app*_ value of 4.6 × 10^-6^ cm/s, indicative of efficient passive diffusion across lipid membranes. LT12 displayed strong chemical stability under simulated gastrointestinal conditions, retaining >92% integrity in simulated gastric fluid (pH 1.2) and >90% in simulated intestinal fluid (pH 6.8) after 2 hours of incubation. In vitro metabolic stability studies further demonstrated low intrinsic clearance. In mouse liver microsomes, **LT12** exhibited a half-life of 118.4 ± 6.7 min (CL_int_ = 15.8 ± 1.3 µL/min/mg), while in human liver microsomes, a longer half-life of 132.6 ± 7.9 min (CL_int_ = 12.7 ± 1.1 µL/min/mg) was observed. Consistent with its chemical stability, **LT12** showed excellent stability in both mouse and human plasma (t1/2 >180 min) and remained stable in PBS for over 5 hours (t1/2 >300 min), suggesting minimal susceptibility to hydrolytic or enzymatic degradation. Plasma protein binding was moderately high (94.2 ± 0.61%), indicating substantial but not excessive serum protein association, which may contribute to sustained systemic exposure while preserving a measurable free fraction.

In safety liability profiling (Table 1), **LT12** showed no cytotoxicity in normal human fibroblast cell lines (WI-38 and HS-27) at concentrations up to 100 µM. Additionally, LT12 exhibited weak inhibition of the hERG potassium channel (IC_50_>100 µM), indicating a low risk of cardiotoxicity. Evaluation against major cytochrome P450 isoforms revealed minimal inhibition at 10 µM, including CYP3A4 (5.9%), CYP2D6 (2.8%), CYP2C9 (4.7%), and CYP2C19 (3.5%), suggesting a low potential for CYP-mediated drug-drug interactions. Taken together, these data demonstrate that **LT12** combines favorable solubility, high membrane permeability, metabolic stability, and a clean early safety profile. Importantly, its physicochemical and permeability characteristics are consistent with a profile suitable for CNS drug discovery.

To evaluate the translational potential of **LT12**, in vivo PK studies were conducted in C57BL/6 mice following single-dose intravenous (IV) and oral administration. After IV dosing at 2 mg/kg, **LT12** exhibited a plasma half-life (t1/2) of 3.6 h, a systemic clearance of 9.8 mL/min/kg, and a volume of distribution (Vd) of 3.1 L/kg, consistent with moderate clearance and broad tissue distribution. The corresponding plasma exposure (AUC0–∞) was 9.9 µM·h, in line with its favorable in vitro metabolic stability.

Following oral administration at 25 mg/kg, **LT12** was efficiently absorbed, reaching a plasma C_max_ of 5.8 µM at a T_max_ of 1.1 h. The total systemic exposure (AUC0–∞) was 58.6 µM·h, corresponding to an oral bioavailability of approximately 47%. The observed plasma half-life of 4.1 h supports sustained exposure over the dosing interval and is consistent with its balanced solubility and permeability profile. Brain exposure was evaluated in parallel given the intended CNS application. **LT12** readily penetrated the brain, achieving a brain C_max_ of 4.7 µM at 1.5 h post-dose and a brain AUC0–∞ of 46.9 µM·h, corresponding to a brain-to-plasma ratio (K_p_) of ~0.8. The similar elimination half-lives in plasma and brain (4.1 h vs 4.4 h) suggest efficient equilibration across the blood-brain barrier without evidence of rapid efflux. The magnitude and duration of CNS exposure indicate that **LT12** maintains pharmacologically relevant concentrations in the brain following oral dosing.

### In Vivo Evaluation Using a Mouse Model of AD

To confirm cross-species target engagement, **LT12** was evaluated against the recombinant extracellular domain of murine ILT3 by MST and exhibited high-affinity binding (Kd = 62.7 ± 5.4 nM, Figure S9), comparable to that observed for the human protein. Thus, **LT12** was advanced for in vivo evaluation in the 5xFAD transgenic mouse model of AD, which recapitulates key features of amyloid pathology, neuroinflammation, and cognitive impairment. Male and female 5xFAD mice and wild-type (WT) littermates (4 months of age) were randomized into treatment groups and administered **LT12** (25 mg/kg, oral, once daily) or vehicle for 21 days (Figure 8A). This time point was selected as it represents a stage of established amyloid deposition and neuroinflammatory activation while remaining responsive to therapeutic intervention.^36^ Behavioral assessment was conducted during days 18-21 of treatment, followed by tissue collection for biochemical and immunological analyses.

**Figure 8.**
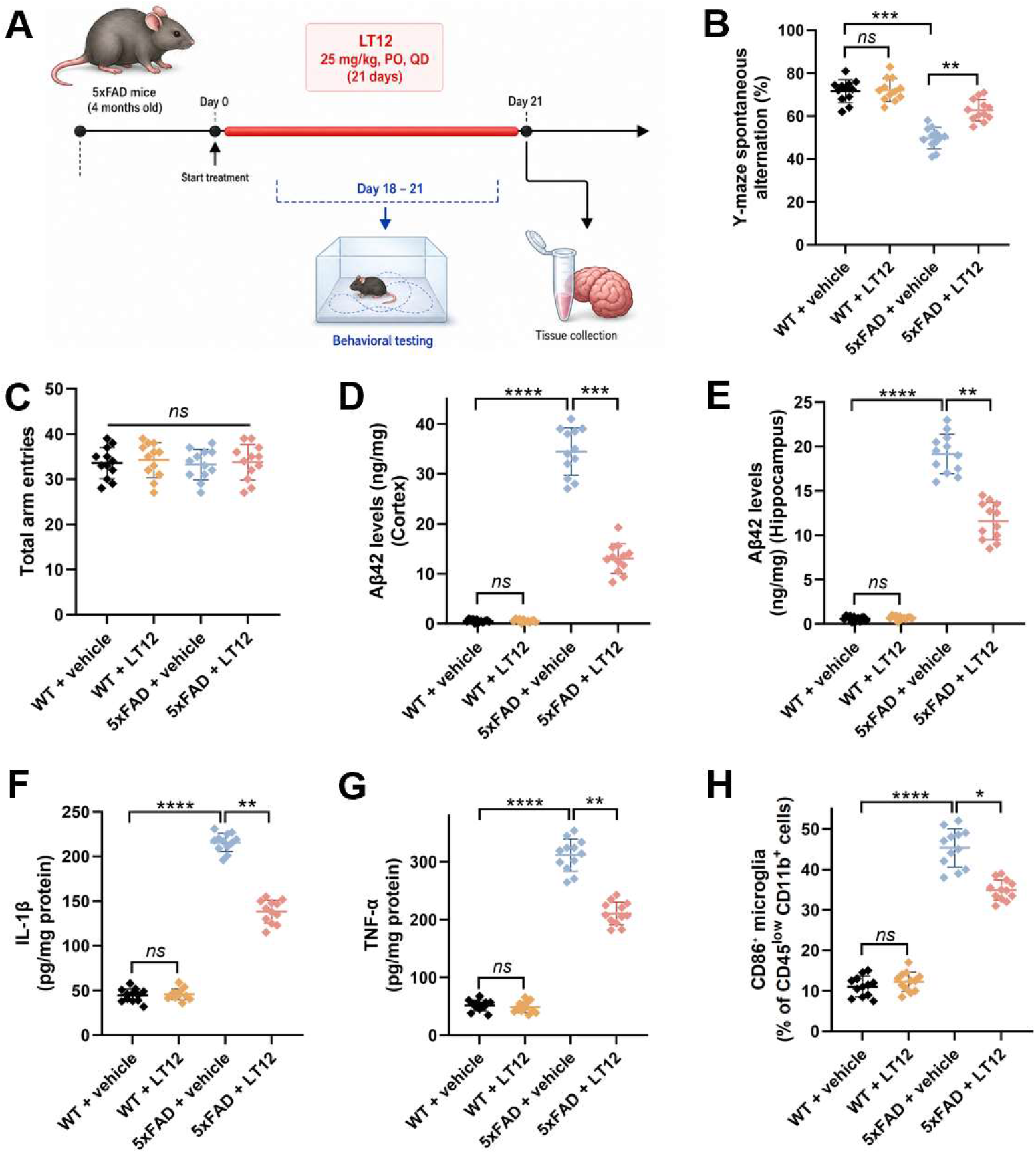
LT12 improves behavioral performance, reduces amyloid burden, and attenuates neuroinflammation in 5xFAD mice. **(A)** Experimental design. Four-month-old 5xFAD mice were treated with **LT12** (25 mg/kg, oral, once daily) for 21 days. **(B)** Y-maze spontaneous alternation (%). 5xFAD mice exhibited impaired performance compared to WT controls, which was improved upon **LT12** treatment. **(C)** Total arm entries. No significant differences were observed across groups. **(D**,**E)** Aβ_42_ levels in cortex **(D)** and hippocampus **(E)**. 5xFAD mice showed elevated Aβ_42_ levels relative to WT, which were significantly reduced following **LT12** treatment. **(F**,**G)** Pro-inflammatory cytokines IL-1β **(F)** and TNF-α **(G)**. Cytokine levels were increased in 5xFAD mice and were significantly decreased upon **LT12** treatment. **(H)** CD86^+^ microglia (% of CD45^low^ CD11b^+^ cells). Microglial activation was elevated in 5xFAD mice and reduced following **LT12** treatment. Data are presented as mean ± SD. Statistical significance was determined by one-way ANOVA with appropriate multiple-comparisons tests. *ns*, not significant; * *p* < 0.05; ** *p* < 0.01; *** *p* < 0.001; **** *p* < 0.0001.

Cognitive performance was evaluated using the Y-maze spontaneous alternation assay (Figure 8B). Vehicle-treated 5xFAD mice displayed a significant reduction in alternation behavior relative to WT controls, consistent with impaired working memory. **LT12** treatment significantly improved performance in 5xFAD mice, restoring alternation levels toward those observed in WT animals, while having no detectable effect in WT controls. Total arm entries were comparable across all groups (Figure 8C), indicating that the observed behavioral improvements were not attributable to changes in locomotor activity.

To determine whether these functional effects were associated with modulation of amyloid pathology, Aβ_42_ levels were quantified in both cortex and hippocampus. As expected, 5xFAD mice exhibited elevated Aβ_42_ levels in both regions compared to WT controls (Figure 8D,E). **LT12** treatment resulted in a significant reduction in Aβ_42_ levels in both cortex and hippocampus, indicating attenuation of amyloid burden. We next assessed neuroinflammatory signaling by measuring levels of pro-inflammatory cytokines. Vehicle-treated 5xFAD mice showed increased expression of IL-1β and TNF-α relative to WT controls (Figure 8F,G). **LT12** treatment significantly reduced both cytokines, consistent with suppression of the inflammatory response in vivo.

Finally, microglial activation was evaluated by quantifying CD86^+^ microglia as a marker of a pro-inflammatory phenotype. 5xFAD mice displayed a marked increase in CD86^+^ microglia compared to WT animals, whereas **LT12** treatment significantly reduced this population (Figure 8H), indicating a shift toward a less activated microglial state. Collectively, these findings demonstrate that **LT12** improves cognitive performance, reduces amyloid burden, and attenuates neuroinflammatory signaling in the 5xFAD model, without affecting general locomotor activity. These results establish a direct link between target engagement and functional efficacy in vivo and support ILT3 modulation as a promising strategy for addressing neuroimmune dysfunction in AD.

## CONCLUSIONS

In this study, we establish ILT3 (LILRB4) as a tractable small molecule target for modulating neuroimmune dysfunction in AD. Using an AS-MS platform, we identified first-in-class small molecule binders that directly engage ILT3, with **LT12** exhibiting nanomolar affinity and robust target engagement across orthogonal biophysical assays. Computational modeling combined with site-directed mutagenesis defined a discrete binding region and revealed distinct interaction modes underlying ligand recognition.

Functionally, pharmacological targeting of ILT3 disrupted the ILT3-ApoE interaction and translated into consistent modulation of downstream signaling and microglial activity. In human iPSC-derived microglia, **LT12** attenuated SHP1/2 phosphorylation, suppressed NF-κB activation, reduced pro-inflammatory cytokine release, and restored Aβ uptake, linking receptor engagement to functional rescue. Importantly, these effects extended in vivo, where **LT12** improved cognitive performance, reduced amyloid burden, and attenuated neuroinflammatory responses in the 5xFAD model. Collectively, these findings provide a comprehensive validation of ILT3 as a druggable neuroimmune checkpoint and highlight small molecule modulation of ILT3 as a promising therapeutic strategy for AD.

## EXPERIMENTAL

### AS-MS Screening and LC-MS Analysis

Recombinant human His-tagged ILT3 protein in PBS (pH 7.4) was used in the AS-MS screening. For all experiments, the protein was freshly thawed and buffer-exchanged into ultrapure water for storage or 50 mM KH2PO4, pH 7.4, with 100 mM NaCl for incubation and screening steps.

The screening collection comprised 5,441 compounds from the Enamine Library, initially evaluated as pooled mixtures containing 1,100 compounds each. In the primary ASMS screening stage, each compound was present at a final concentration of 1 µM, alongside 1 µM ILT3 protein. Incubations were conducted in 50 mM KH2PO4 buffer (pH 7.4) supplemented with 100 mM NaCl. To maintain weak and transient binding interactions, samples were incubated at 4 °C under low-agitation conditions. For hit validation, selected pools were deconvoluted and reconstituted into smaller subsets of approximately 8 compounds per mixture. These were reassessed using an increased ILT3 concentration of 2.5 µM while keeping compound concentrations constant at 1 µM. This secondary format improved sensitivity for reproducible binders and reduced potential masking effects arising from competition within larger pools.

ASMS analysis was performed using a two-dimensional SEC-LC-MS workflow. Following incubation, protein-ligand mixtures were injected directly onto a PolyLC size-exclusion column (2.1 × 50 mm, 5 µm) maintained at 8 °C. Separation was carried out using either PBS or 50 mM KH2PO4 buffer (pH 7.4) at a flow rate of 1.0 mL/min. High-molecular-weight fractions corresponding to protein-ligand complexes were collected online and transferred to a second-dimension reversed-phase LC-MS system. In the second dimension, analytes were separated on an Agilent RRHD C18 column (2.1 × 50 mm, 1.7 µm) held at 60 °C. The mobile phases consisted of 0.2% formic acid in water containing 50 mM ammonium acetate (aqueous phase) and 0.2% formic acid in acetonitrile (organic phase). The gradient program included an initial hold at 2% organic solvent (0-0.5 min), followed by a linear increase to 95% organic (0.5-2.5 min), a hold at 95% (up to 3.5 min), rapid re-equilibration to 2% (by 3.6 min), and final equilibration until 5.0 min. The flow rate was set to 0.3 mL/min with an injection volume of 40 µL. This orthogonal separation strategy enabled efficient desalting, improved chromatographic resolution, and optimal compatibility with downstream mass spectrometric detection.

### Hit Identification Criteria

Mass spectrometric data were acquired in positive ionization mode using a high-resolution time-of-flight (TOF) instrument calibrated to achieve mass accuracy within 5 ppm. Signals detected in protein-containing samples (P^+^) were systematically compared against matched protein-free controls (P^−^) to assess enrichment. Compounds were classified as hits based on the following criteria: (i) measured monoisotopic mass within ±5 ppm of the expected value, (ii) consistent chromatographic retention time across replicate P^+^ analyses, (iii) a well-defined and reproducible isotopic distribution, and (iv) markedly diminished or undetectable signal in the corresponding P^−^ control. Signal intensities were averaged across replicates to improve confidence in hit assignment. Application of these criteria to the primary screen yielded 36 candidate ILT3 binders from a total of 5,441 compounds. Subsequent retesting in reduced pool formats under elevated protein concentration (2.5 μM of ILT3) confirmed 18 compounds as reproducible binders.

### Dianthus TRIC Assay

Initial screening for small molecule ILT3 binders was performed using the temperature-related intensity change (TRIC) assay on a Dianthus NT.23 Pico instrument (NanoTemper Technologies). Recombinant human His-tagged ILT3 protein was fluorescently labeled with RED-tris-NTA 2nd Generation dye (NanoTemper, Cat. #MO-L018) according to the manufacturer’s protocol.

Labeled ILT3 (final concentration: 30 nM) was incubated with test compounds at a fixed concentration (50 µM) in MST buffer (PBS pH 7.4 supplemented with 0.05% Tween-20 and 1% (v/v) DMSO). Fluorescence was measured in Dianthus 384-well plates (NanoTemper Technologies, Catalog# DI-P001A) with LED power set to 40%. Fluorescence was recorded at 670 nm and 650 nm, and normalized fluorescence (F_norm_) was calculated as the ratio of F670/F650. Binding responses were quantified as Fnorm. Compounds with ΔF_norm_ ≥ 5 units and reproducible responses across replicates (n = 5) were classified as hits.

### Quantitative Binding Affinity Determination Using Monolith

Binding affinities of selected hits were quantified by microscale thermophoresis using a Monolith X instrument (NanoTemper Technologies). His-tagged ILT3 protein was labeled with RED-tris-NTA 2nd Generation dye using the Monolith His-Tag Labeling Kit (Cat. #MO-L018) following the manufacturer’s guidelines. Compound titrations were prepared as serial dilutions (PBS buffer, pH 7.4, 0.05% Tween-20, 1% (v/v) DMSO). Following a 15 min incubation at room temperature in the dark, samples were loaded into Monolith capillaries (Cat. #MO-K022) and analyzed at 25 °C using 40% LED power and medium MST power settings. Normalized fluorescence (F_norm_) values were determined as the ratio of fluorescence intensity after and before IR laser heating. Each compound was evaluated in five technical replicates. Dissociation constants (Kd) were calculated from three independent experiments using MO.Affinity Analysis software and GraphPad Prism 10, applying standard dose-response fitting models.

### SPR analysis

SPR validation of ILT3 binding was performed using Biacore™ 8K SPR system (Cytiva, Marlborough, MA, USA). ILT3-His Protein (10 μg/mL in PBS, pH 7.4) was immobilized on a Series S Sensor Chip CM5 (29104988, Cytiva, Marlborough, MA, USA) using a commercial amine coupling kit (BR100050, Cytiva, Marlborough, MA, USA). Immobilization was performed at a flow rate of 10 μL/min for 420 s, followed by blocking with ethanolamine. The flow cell with solely ethanolamine-block on the same channel served as the reference. Immobilization buffer: PBS-P+ (28995084, Cytiva, Marlborough, MA, USA). Briefly, gradient concentrations of the compound were prepared in the assay buffer, and injected over the sensor chip in a single-cycle kinetics model with a flow rate at 30 μL/min for 120 s per injection. After each injection, a 30 s-regeneration was performed at a flow rate of 30 μL/min using the regeneration buffer. Assays were performed in triplicate, and Kd values are reported as mean ± SD.

### Computational studies

Molecular docking, molecular dynamics (MD) simulations, and binding free energy calculations were performed to characterize the interactions of **LT6** and **LT12** with ILT3. The crystal structure of ILT3 (PDB ID: 3P2T) was obtained from the Protein Data Bank. The solvents and water were removed from the receptor prior to docking. The lead compounds in SMILES format were converted into .mol2 and subsequently to .pdbqt using openbabel (37) and MGLTools (38). The pdbqt file includes Gasteiger charges and information on rotatable bonds. **LT6** is positively charged, and **LT12** is neutral. We have carried out molecular docking using AutoDock4.0 (39). The top binding pose with the lowest docking energy was selected as the input for the subsequent MD simulations. Both ligands bind to different binding sites in ILT3 which is to maximize the complementary interactions arising from hydrophobic, electrostatic and hydrogen bonding interactions between the receptor and ligand atoms.

The .frcmod and .prep files for both ligands were prepared using the Antechamber package of AmberTools. The complex structures of both ligands bound to the ILT3 receptor were prepared using the best-binding pose obtained from molecular docking. The minimization run was performed on these complexes, followed by simulations in the constant-volume and isothermal-isobaric ensembles. The simulations were carried out at ambient temperature and pressure. The molecular dynamics simulations were carried out using Amber22 software. The time step for solving the equation of motion was chosen as 2 fs. The equilibration runs were carried out for 10 ns, and the production runs for 80 ns. Various analyses, including root-mean-square displacement, radius of gyration, and hydrogen-bond analysis, were performed on configurations from the production runs. Further free energy calculations for estimating the binding affinity of the ligands for the ILT3 complex were performed by employing the molecular mechanics-Generalized Born approach (40). The free energy of binding is obtained as the difference between the free energy of the complex and the sum of free energies of the receptor and the ligand. The free energy calculations use an implicit solvent approximation, so receptor, ligand, and complex structures from MD are used only as inputs. The free energies are obtained as the sum of van der Waals, electrostatic, and solvation energies. The solvation energies include the contributions from both polar and non-polar components. We have also carried out residue-wise decomposition analysis of free energies.

### HPLC Purity

The purity of **LT12** was confirmed to be >99% using HPLC (Figure S10). HPLC analysis was performed with a reversed-phase column Phenomenex Gemini, C18 (250 mm × 4.60 mm, 5 μm) on an HPLC Agilent system. The mobile phase used was an acetonitrile-H_2_O gradient and a 1 mL/min flow rate. UV absorption at 215 and 254 nm was used to monitor the method.

### ELISA-based competition assay

ILT3 extracellular domain (2 µg/mL) was immobilized on 96-well high-binding plates (Corning) overnight at 4°C. Plates were blocked with 5% BSA in PBS for 1 h at room temperature. Recombinant human ApoE (R&D Systems) was added at a fixed concentration (100 nM) in the presence of serial dilutions of compounds. Bound ApoE was detected using an HRP-conjugated anti-ApoE antibody, followed by TMB substrate development. Absorbance was measured at 450 nm using a Tecan plate reader. Data were normalized to DMSO controls and fitted to determine IC_50_ values using nonlinear regression (GraphPad Prism v10). Reported values represent mean ± SD (n=5).

### BLI competition study

BLI experiments were performed using an Octet RED96 system (Sartorius). LILRB4 protein was immobilized onto Ni-NTA biosensors, followed by equilibration in assay buffer (PBS + 0.05% Tween-20). ApoE (100 nM) was associated in the presence of increasing concentrations of compounds. Binding responses were monitored as wavelength shifts, and inhibition curves were generated to determine IC_50_ values. Data were analyzed using Octet Data Analysis software. Reported values represent mean ± SD (n=5).

### Site-directed mutagenesis

Alanine substitutions (N20A, V15A, P12A, I16A, P89A, and E91A) were introduced into the human ILT3 extracellular domain using the QuikChange Lightning Site-Directed Mutagenesis Kit (Agilent Technologies) according to the manufacturer’s protocol. Briefly, mutagenic primers were designed to incorporate the desired codon substitutions and PCR amplification was performed using a high-fidelity polymerase. Parental methylated DNA was digested with DpnI, and the resulting plasmids were transformed into *E. coli* DH5α cells. All mutations were confirmed by Sanger sequencing. Recombinant wild-type and mutant ILT3 proteins were expressed and purified as described above. Protein integrity and purity (>90%) were verified by SDS-PAGE and analytical size-exclusion chromatography. Binding affinities of mutant proteins were evaluated by MST under identical conditions to wild-type controls, and Kd values were obtained from nonlinear regression fits as described above.

### Human iPSC-derived microglia assays

Human iPSC-derived microglia (FujiFilm Cellular Dynamics) were cultured in microglia maintenance medium (Axol Bioscience Ltd, Catalog #ax0660) supplemented according to the supplier’s recommendations at 37 °C in a humidified 5% CO_2_ incubator. Cells were plated at 50,000 cells per well in poly-D-lysine-coated 96-well plates and allowed to recover for 24 h prior to treatment. Aβ42 oligomers were prepared by dissolving lyophilized Aβ42 peptide (≥95% purity; AnaSpec, Catalog# AS-20276) in HFIP, followed by evaporation, resuspension in DMSO, and oligomerization in PBS at 4 °C for 24 h. Cells were treated with Aβ42 (1 μM) in the presence or absence of recombinant human ApoE (100 nM), followed by addition of **LT12** at indicated concentrations (0.5-5 μM). Treatments were performed for 24 h. Vehicle controls contained equivalent DMSO concentrations (<0.5%).

### Signaling assays

Phosphorylation of SHP1 and SHP2 was quantified using phospho-specific ELISA kits (abcam, catalog# ab279924 and ab314344, respectively) following the manufacturer’s protocol. Briefly, cells were lysed in ice-cold lysis buffer supplemented with protease and phosphatase inhibitors. Lysates were normalized for total protein content (BCA assay), and equal amounts were loaded per well. Reported values represent mean ± SD (n=5).

NF-κB activation was measured using a luciferase reporter assay in human iPSC-derived microglia. Cells were plated at 50,000 cells per well in white 96-well plates and transfected with an NF-κB firefly luciferase reporter (Promega) and a Renilla luciferase control plasmid (Promega) using Lipofectamine 3000 (Thermo Fisher Scientific, Cat. #L3000008). Twenty-four hours post-transfection, cells were treated with Aβ42 oligomers (1 μM) ± ApoE (100 nM) and **LT12** (0.5-5 μM) for 24 h. Luciferase activity was measured using the Dual-Glo Luciferase Assay System (Promega, Cat. #E2920). NF-κB activity was calculated as the ratio of firefly to Renilla signal and expressed as fold change relative to vehicle controls. Reported values represent mean ± SD (n=5).

### Cytokine measurements

IL-1β levels in culture supernatants were quantified using a human IL-1β ELISA kit (R&D Systems, catalog# DLB50). Supernatants were collected, clarified by centrifugation (1,000 × g, 5 min), and analyzed according to the manufacturer’s instructions. Cytokine concentrations were interpolated from standard curves and normalized where appropriate. Reported values represent mean ± SD (n=5).

### Aβ uptake assay

Fluorescently labeled Aβ42 (HiLyte Fluor 555-conjugated from AnaSpec, catalog# AS-60480-01) was added to cells at a final concentration of 500 nM following treatment conditions described above. After incubation (4 h), cells were washed extensively with PBS and treated with trypan blue to quench extracellular fluorescence. Intracellular fluorescence was quantified using a Tecan Spark plate reader and normalized to cell number (total protein).

### Cell viability

Cell viability was assessed using the CellTiter-Glo luminescent assay (Promega, catalog#G7570) according to the manufacturer’s protocol. Following treatment, reagent was added directly to wells, incubated for 10 min at room temperature, and luminescence was measured. Viability was normalized to vehicle-treated controls. Reported values represent mean ± SD (n=5).

### PK studies

In vitro PK profiling was performed as we previously reported.^13^ In vivo PK studies were conducted in male C57BL/6 mice (8-10 weeks old; n = 5 per time point). **LT12** was administered by intravenous injection (2 mg/kg) or oral gavage (25 mg/kg) in a formulation consisting of 5% DMSO, 40% PEG400, and 55% sterile saline (v/v/v), prepared fresh prior to dosing. Blood samples were collected at 0.25, 0.5, 1, 2, 4, 8, and 24 h post-dose via tail vein sampling into EDTA-coated tubes. Plasma was isolated by centrifugation at 3,000 × g for 10 min at 4 °C.

For CNS exposure analysis, mice were perfused with ice-cold PBS before brain collection to minimize residual blood contamination. Brain tissues were harvested, weighed, and homogenized in ice-cold PBS. Compound concentrations in plasma and brain homogenates were quantified by LC-MS/MS following protein precipitation with acetonitrile containing an internal standard. Chromatographic separation was performed on a C18 column, and analytes were detected in positive electrospray ionization mode using multiple reaction monitoring. Calibration curves were prepared in blank mouse plasma or brain homogenate matrix and used to calculate compound concentrations. PK parameters were calculated by noncompartmental analysis using Phoenix WinNonlin. Brain-to-plasma exposure ratios were calculated from AUC values. Unbound brain-to-plasma ratios (Kp,uu) were calculated using measured plasma and brain unbound fractions.

### In vivo efficacy in 5xFAD mice

Male and female 5xFAD mice and wild-type littermates (4 months old) were randomized into treatment groups (n = 12 per group; 6 males and 6 females per group). **LT12** was administered by oral gavage at 25 mg/kg once daily (QD) for 21 days. Vehicle-treated animals received formulation only. Treatment allocation, behavioral testing, biochemical assays, flow cytometry analysis, and data quantification were performed by investigators blinded to genotype/treatment.

### Behavioral testing

Cognitive function was assessed using the Y-maze spontaneous alternation assay. Mice were placed in the maze and allowed to explore freely for 8 min. Alternation percentage was calculated as the ratio of sequential entries into all three arms over total possible alternations. Total arm entries were recorded to control for locomotor activity.

### Amyloid quantification

Cortex and hippocampus were microdissected separately, snap-frozen, and homogenized in ice-cold extraction buffer containing protease inhibitors. Tissue homogenates were centrifuged to obtain soluble fractions, and the remaining pellets were further extracted to recover insoluble Aβ. Soluble Aβ Was extracted in TBS or PBS-based buffer, and insoluble Aβ was extracted from pellets using 5 M guanidine-HCl or 70% formic acid. Aβ42 levels were quantified using a commercially available ELISA kit (Thermo Fisher Scientific; Aβ42, Cat. # KMB3441), according to the manufacturer’s instructions. Concentrations were interpolated from standard curves and normalized to total protein content determined by BCA assay. Data are reported separately for cortex and hippocampus.

### Cytokine analysis

Brain homogenates were prepared in lysis buffer and cytokine levels (IL-1β, TNF-α) were measured using ELISA kits (R&D Systems, catalog# MLB00C and MTA00B, respectively). Values were normalized to total protein content.

### Microglial activation

Brain tissues (cortex and hippocampus) were rapidly dissected and processed into single-cell suspensions using the Adult Brain Dissociation Kit (Miltenyi Biotec, Cat. #130-107-677) according to the manufacturer’s protocol. Myelin was removed using Myelin Removal Beads II (Miltenyi Biotec, Cat. #130-096-733). Cells were resuspended in FACS buffer (PBS containing 2% fetal bovine serum and 2 mM EDTA) and incubated with anti-mouse CD16/CD32 Fc block (BioLegend, Cat. #101320) for 10 min at 4 °C to prevent non-specific binding.

Cells were stained for 30 min at 4 °C with fluorophore-conjugated antibodies against CD45 (APC, clone 30-F11, BioLegend, Cat. #103112), CD11b (FITC, clone M1/70, BioLegend, Cat. #101206), TMEM119 (PE, clone 106-6, BioLegend, Cat. #157306), and CD86 (PerCP-Cy5.5, clone GL-1, BioLegend, Cat. #105028). Dead cells were excluded using a viability dye (Zombie NIR, BioLegend, Cat. #423106). Data were acquired on a BD LSRFortessa flow cytometer (BD Biosciences) and analyzed using FlowJo v10. Microglia were defined as live CD45^low^ CD11b^+^ TMEM119^+^ cells. Activated microglia were quantified as the percentage of CD86^+^ cells within the microglial population. A minimum of 50,000 live events per sample were recorded.

### Statistical analysis

All data are presented as mean ± SD. Statistical analyses were performed using GraphPad Prism v10. Two-group comparisons were conducted using unpaired two-tailed Student’s t-tests. Multiple comparisons were analyzed using one-way ANOVA followed by Tukey’s post hoc test. *P* values < 0.05 were considered statistically significant. Exact sample sizes and statistical tests are indicated in figure legends.

## Supporting Information

The Supporting Information is available free of charge.

### ABBREVIATIONS

AD: Alzheimer’s disease
ADME: absorption, distribution, metabolism, and excretion
ApoE: apolipoprotein E
AS-MS: affinity selection–mass spectrometry
AUC: area under the curve
Aβ: amyloid-β
Aβ_42_: amyloid-β 42
BLI: biolayer interferometry
CL: clearance
CLint: intrinsic clearance
Cmax: maximum concentration
CNS: central nervous system
DMSO: dimethyl sulfoxide
ELISA: enzyme-linked immunosorbent assay
FaSSIF: fasted-state simulated intestinal fluid
Fnorm: normalized fluorescence
IC_50_: half-maximal inhibitory concentration
IL-1β: interleukin-1 beta
ILT3: immunoglobulin-like transcript 3
iPSC: induced pluripotent stem cell
ITIM: immunoreceptor tyrosine-based inhibitory motif
Kd: equilibrium dissociation constant
Kp: brain-to-plasma concentration ratio
LC-MS: liquid chromatography–mass spectrometry
LILRB4: leukocyte immunoglobulin-like receptor B4
MD: molecular dynamics
MM/GBSA: molecular mechanics/generalized Born surface area
MST: microscale thermophoresis
NF-κB: nuclear factor kappa-light-chain-enhancer of activated B cells
PAMPA: parallel artificial membrane permeability assay
PBS: phosphate-buffered saline
PBST: phosphate-buffered saline with Tween-20
PK: pharmacokinetics
PPB: plasma protein binding
SEC: size-exclusion chromatography
SHP1/2: Src homology region 2 domain-containing phosphatase 1/2
SPR: surface plasmon resonance
t1/2: half-life
Tmax: time to maximum concentration
TNF-α: tumor necrosis factor alpha
TRIC: temperature-related intensity change
Vd: volume of distribution
WT: wild-type.

## Notes

The authors declare no competing financial interests.

