## Supporting Information for "Discovery of ILT3 (LILRB4) Small Molecule Inhibitors by Affinity Selection-Mass Spectrometry Reveals Druggability of a Neuroimmune Checkpoint in Alzheimer’s Disease"

*Electronic Supplementary Information*

| **Contents** | |  |
| --- | --- | --- |
| Chemical structures of the preliminary hit compounds identified by SEC-coupled AS-MS screening | | S2 |
| MST dose-response curve for **LT6** | | S4 |
| 3D Docking pose of the predicted binding mode of **LT6** in ILT3  Two-dimensional interaction map highlighting key predicted interactions of **LT6** with ILT3  The binding free energies of the **LT6** and **LT12** with the ILT3 receptor | | S4  S5  S5 |
| Dose-response curves from ELISA showing inhibition of ApoE binding to immobilized ILT3 in the presence of increasing concentrations of **LT6**  BLI analysis of ApoE binding to ILT3 in the presence of **LT6**  **LT12** suppresses SHP2 phosphorylation in iPSC-derived human microglia  **LT12** does not affect viability of iPSC-derived human microglia  Cross-species binding of **LT12** to murine ILT3 measured by MST  HPLC trace for compound **LT12**. | | S6  S6  S7  S9  S9  S10 |

**Table S1**. Chemical structures of the preliminary hit compounds identified by SEC-coupled AS-MS screening.

| **Compound name** | **Chemical structure** |
| --- | --- |
| **LT1** | 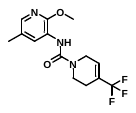 |
| **LT2** | 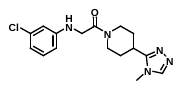 |
| **LT3** | 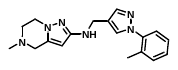 |
| **LT4** | 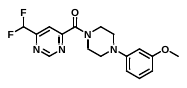 |
| **LT5** | 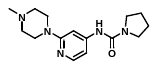 |
| **LT6** | 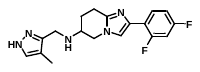 |
| **LT7** | 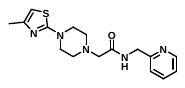 |
| **LT8** | 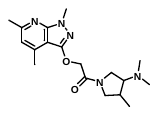 |
| **LT9** | 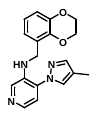 |
| **LT10** | 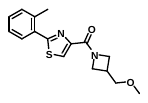 |
| **LT11** | 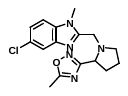 |
| **LT12** | 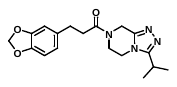 |
| **LT13** | 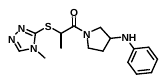 |
| **LT14** | 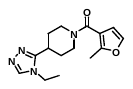 |
| **LT15** | 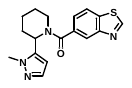 |
| **LT16** | 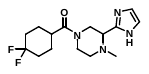 |

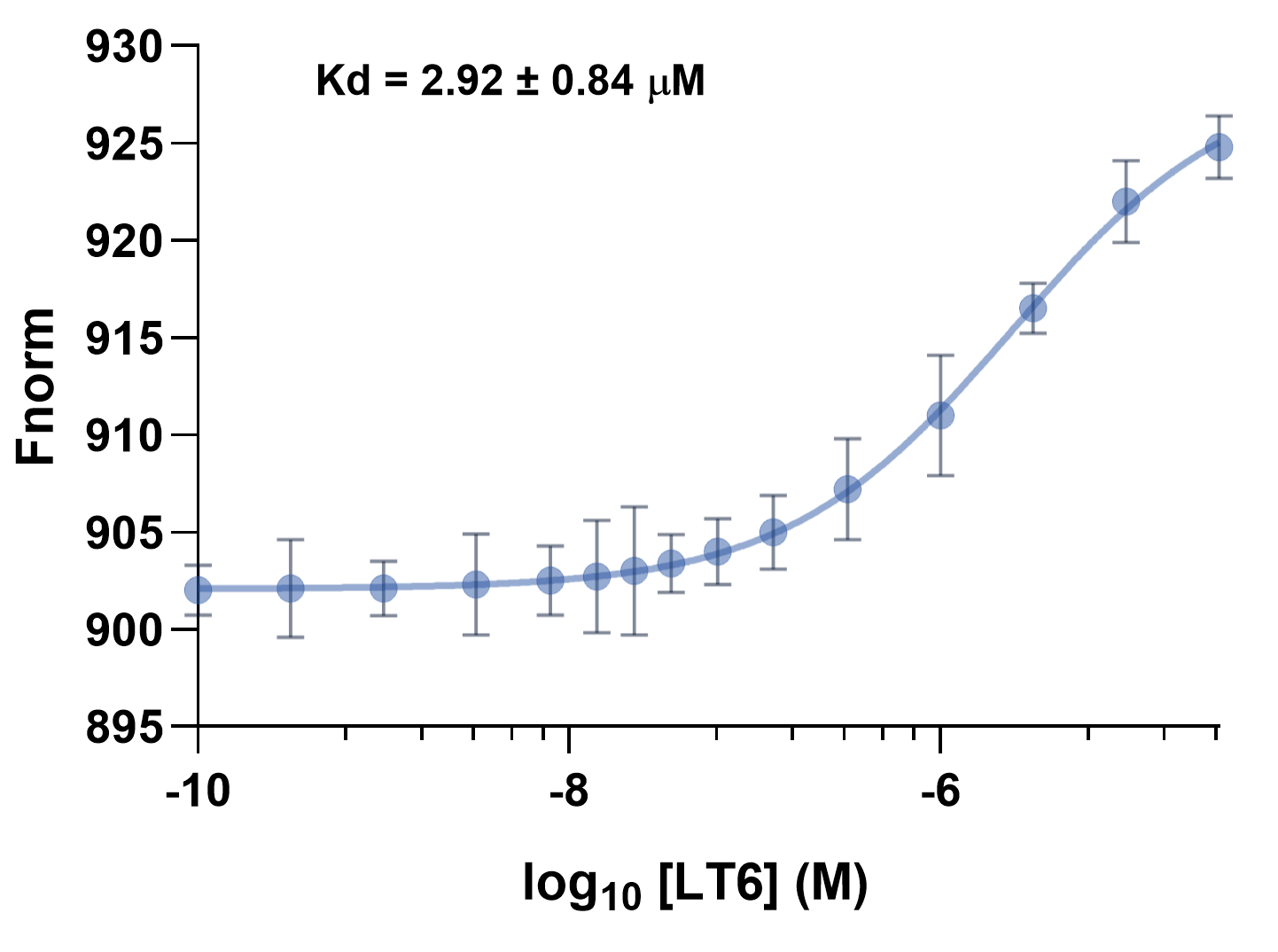

**Figure S1.** MST dose-response curve for **LT6**. Data are shown as mean ± SD (n = 5).

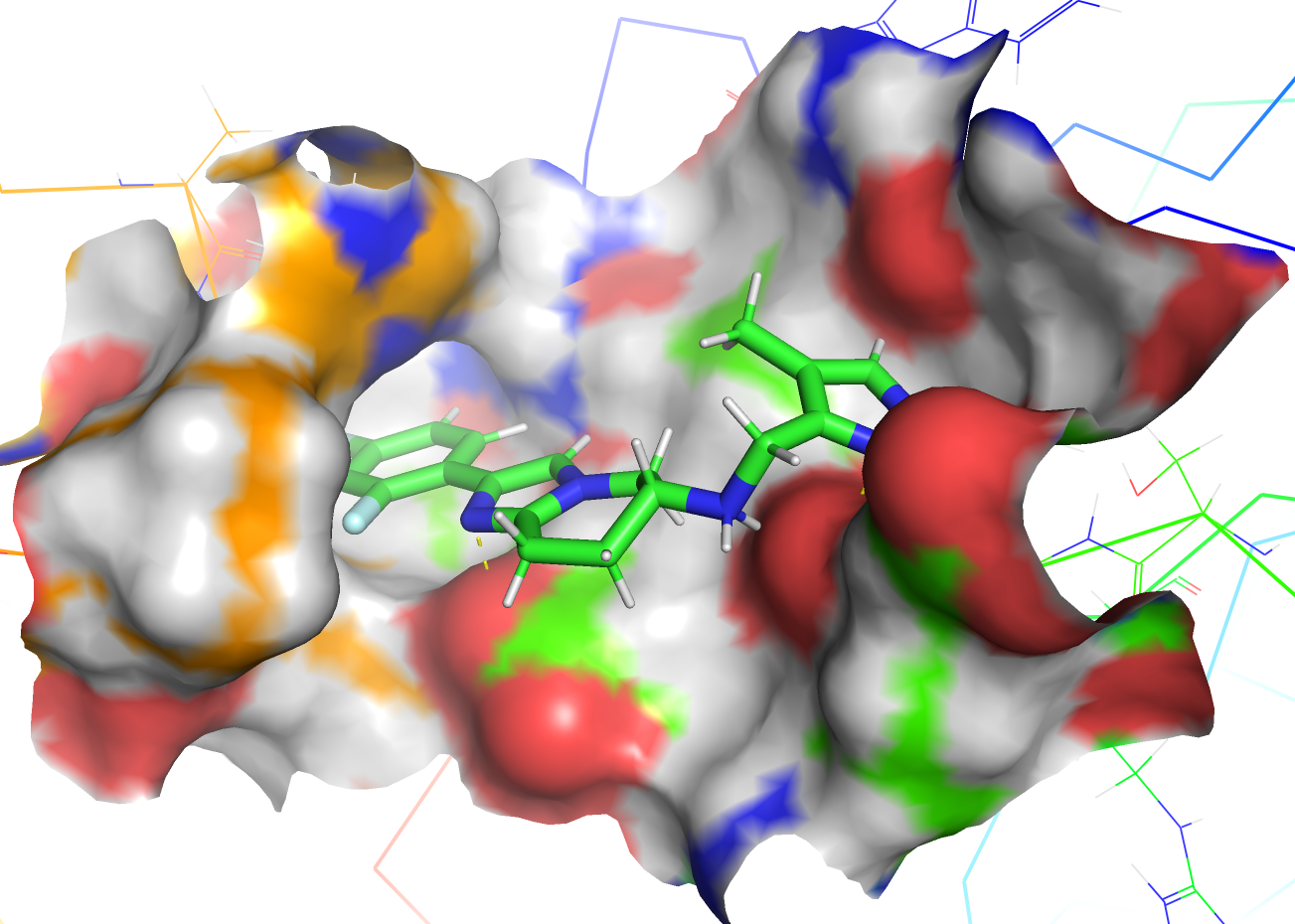

**Figure S2. 3D Docking pose of the predicted binding mode of LT6 in ILT3.** Three-dimensional docked pose of **LT6** (green) within the ILT3 binding pocket, shown on the protein surface colored by physicochemical properties.

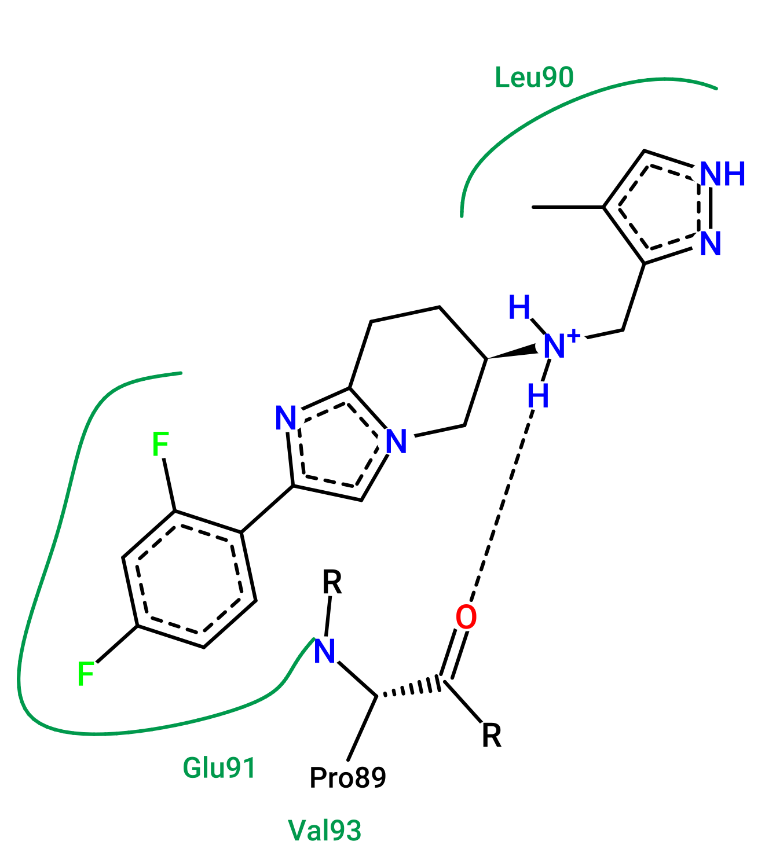

**Figure S3. Two-dimensional interaction map highlighting key predicted interactions of LT6 with ILT3.** Dashed lines represent hydrogen bonds and solid green lines represent hydrophobic interactions.

**Table S2. The binding free energies of the LT6 and LT12 with the ILT3 receptor.**

| System | 𝚫E_VDW, kcal/mol | 𝚫E_ELEC, kcal/mol | 𝚫E_ELEC, kcal/mol | 𝚫E_VDW, kcal/mol | 𝚫G_total, kcal/mol |
| --- | --- | --- | --- | --- | --- |
| **LT6** | -14.3 | -112.4 | 117.3 | -2.0 | -11.4 |
| **LT12** | -31.6 | -10.9 | 21.3 | -3.9 | -25.1 |

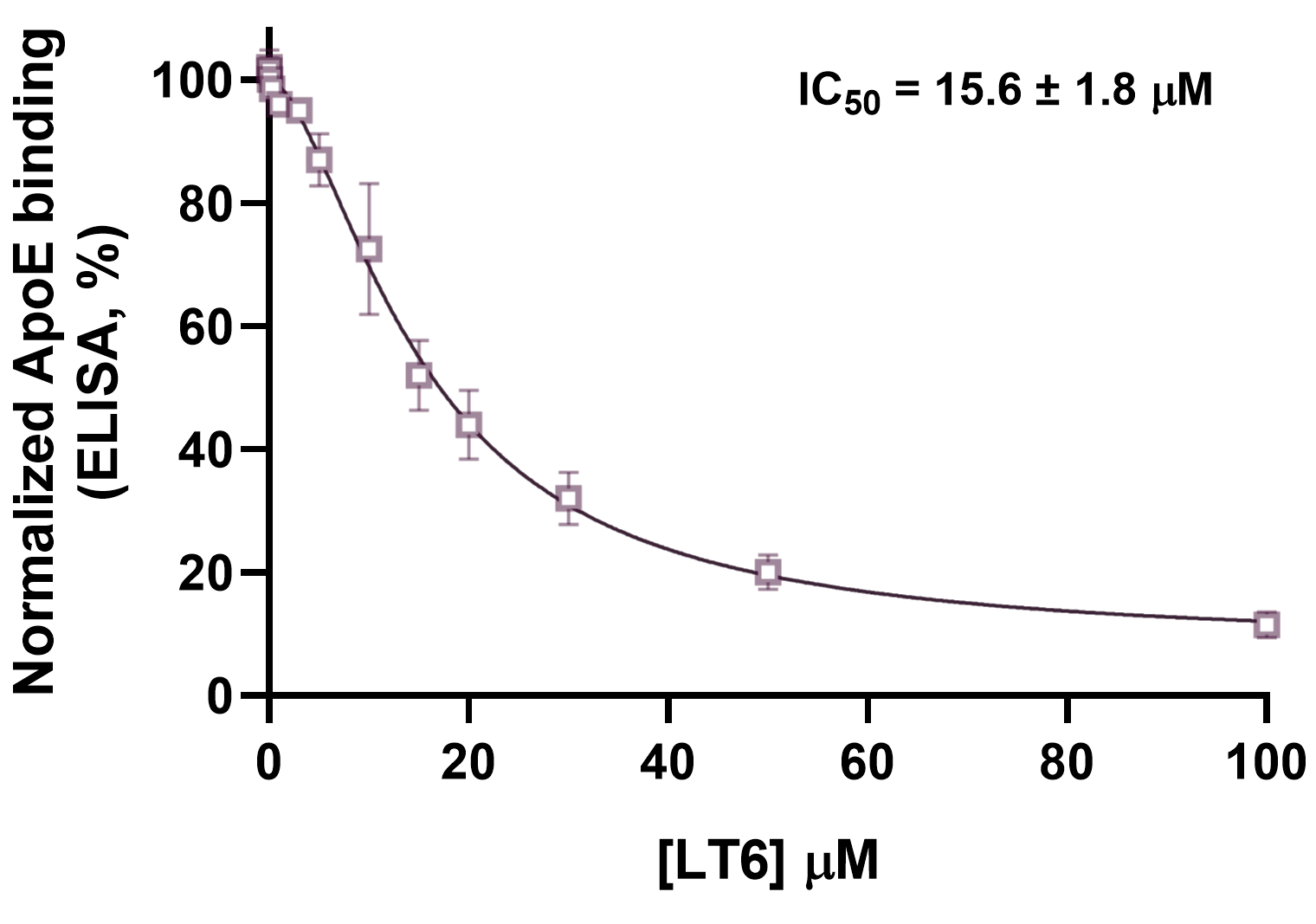

**Figure S4.** Dose-response curves from ELISA showing inhibition of ApoE binding to immobilized ILT3 in the presence of increasing concentrations of **LT6**. Data are presented as mean ± SD (n=5).

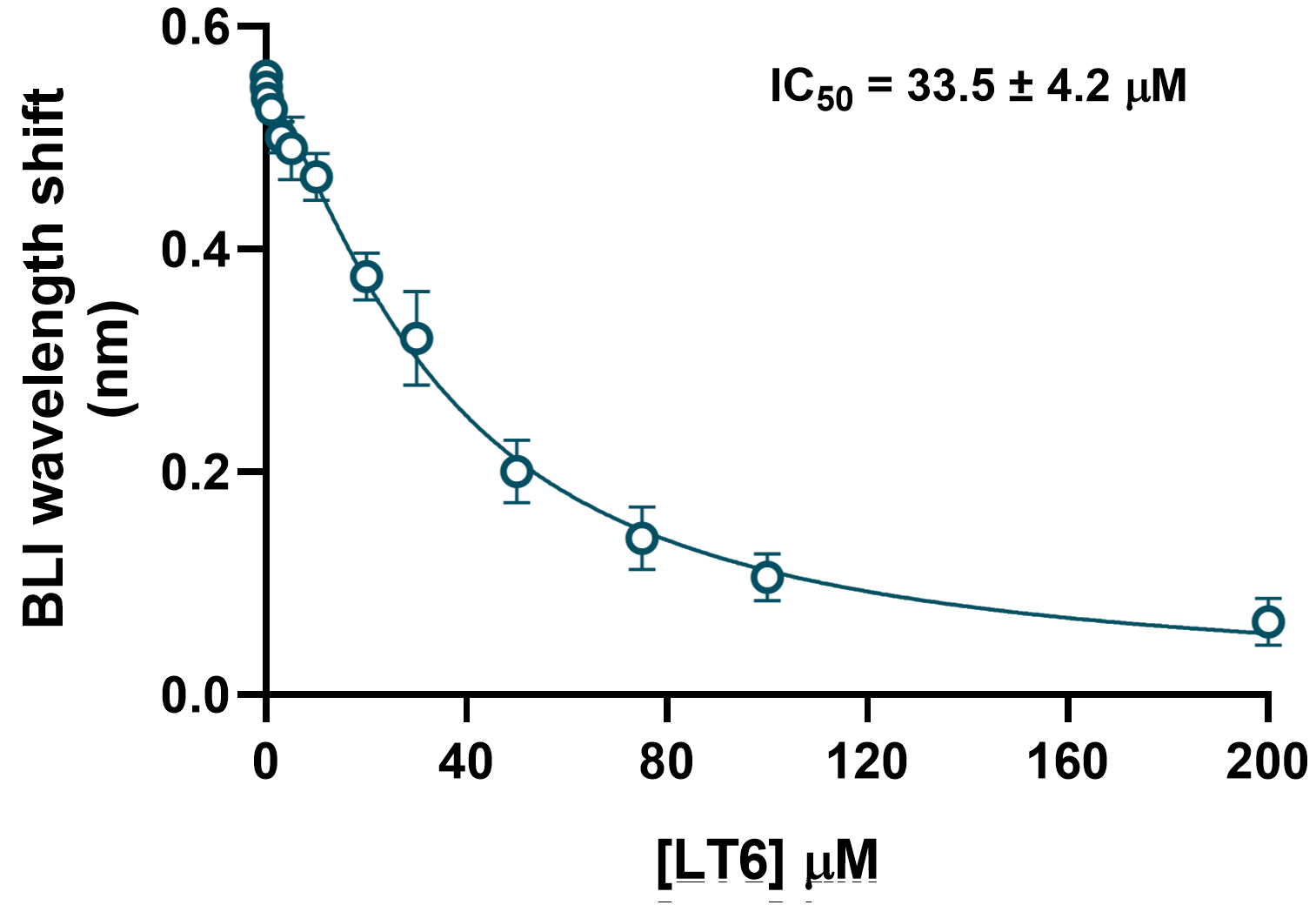

**Figure S5.** BLI analysis of ApoE binding to ILT3 in the presence of **LT6**. Data are presented as mean ± SD (n=5).

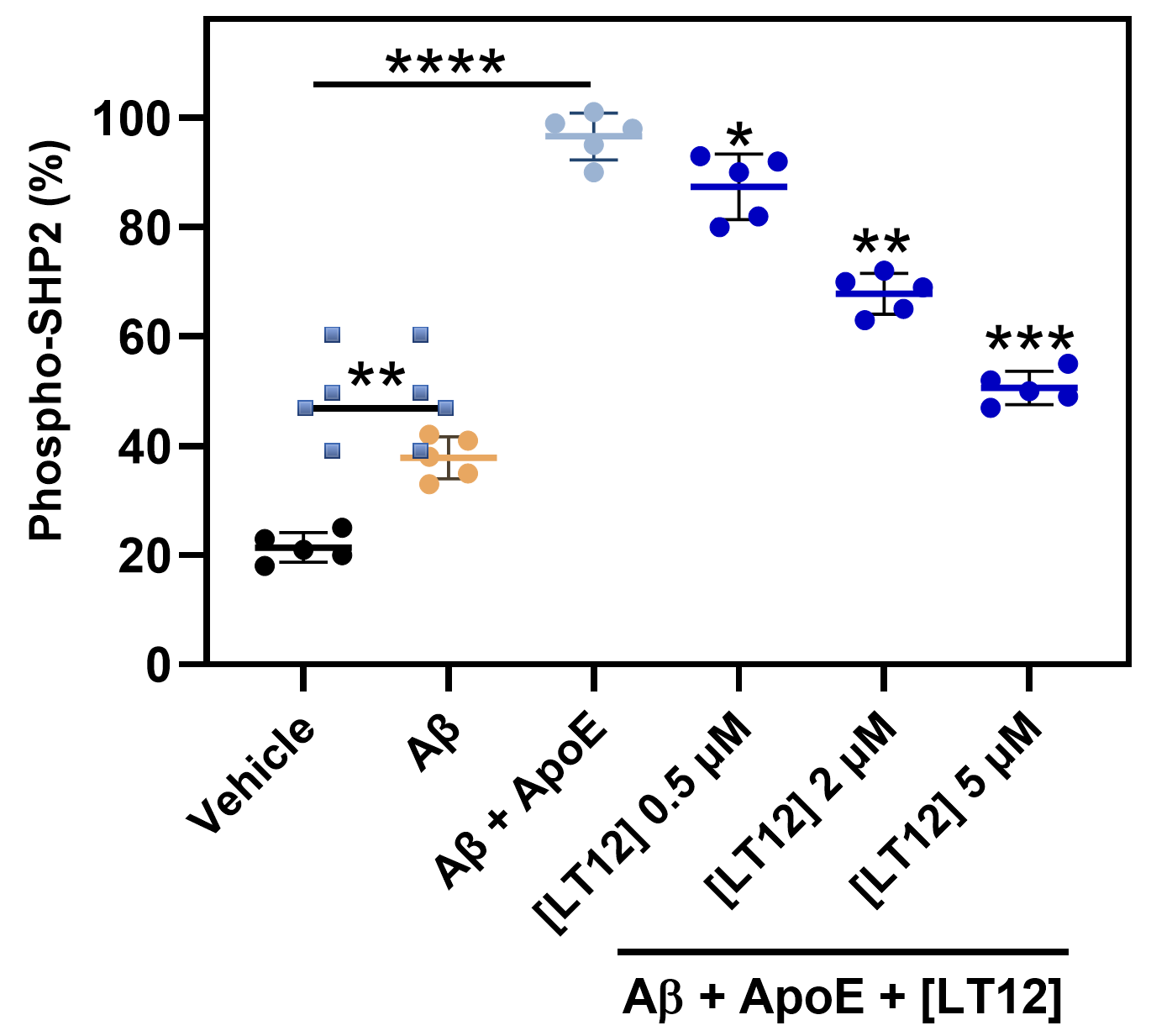

**Figure S6. LT12 suppresses SHP2 phosphorylation in iPSC-derived human microglia.**
Phosphorylation of SHP2 following stimulation with Aβ₄₂ and ApoE. Co-treatment with ApoE increased SHP2 phosphorylation relative to Aβ₄₂ alone, consistent with activation of ILT3-dependent signaling. **LT12** reduced phospho-SHP2 levels in a concentration-dependent manner. Data are presented as mean ± SD (n = 5). Statistical significance was determined by one-way ANOVA with appropriate multiple-comparisons tests. Comparisons between Aβ₄₂ and vehicle, and between Aβ₄₂ + ApoE and control conditions, are indicated where appropriate. For compound-treated groups, statistical comparisons were performed relative to the Aβ₄₂ + ApoE condition unless otherwise stated. **p* < 0.05, ***p* < 0.01, ****p* < 0.001, *****p* < 0.0001.

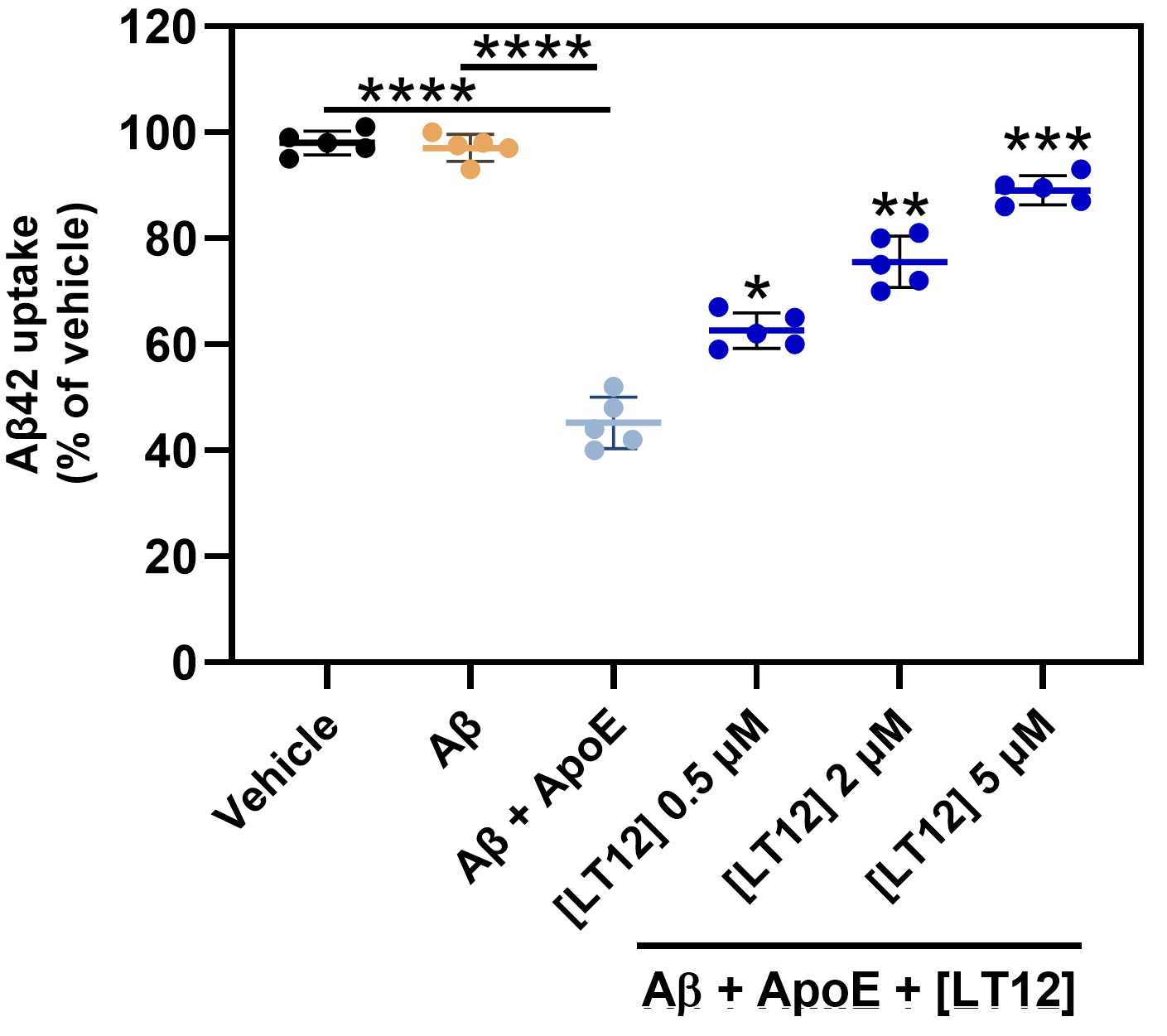

**Figure S7. LT12 restores Aβ₄₂ uptake in iPSC-derived human microglia.**
Aβ₄₂ uptake (% of vehicle) under indicated treatment conditions. Aβ₄₂ alone did not significantly alter uptake relative to vehicle, whereas co-treatment with ApoE reduced Aβ internalization. **LT12** treatment increased Aβ uptake in a concentration-dependent manner. Data are presented as mean ± SD (n = 5). Statistical significance was determined by one-way ANOVA with appropriate multiple-comparisons tests. Comparisons between Aβ₄₂ and vehicle, and between Aβ₄₂ + ApoE and control conditions, are indicated where appropriate. For compound-treated groups, statistical comparisons were performed relative to the Aβ₄₂ + ApoE condition unless otherwise stated. **p* < 0.05, ***p* < 0.01, ****p* < 0.001, *****p* < 0.0001.

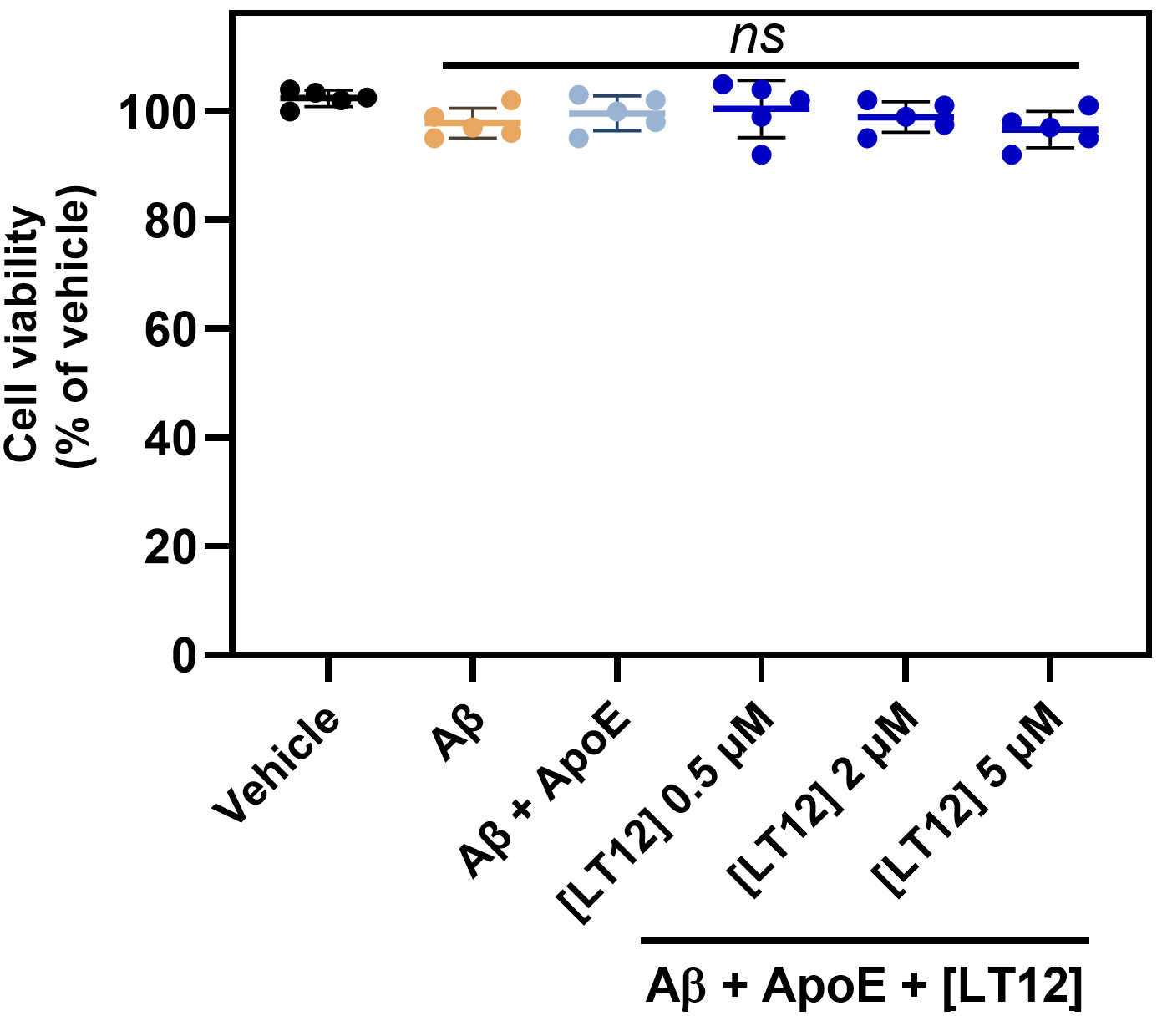

**Figure S8. LT12 does not affect viability of iPSC-derived human microglia.**
Cell viability (% of vehicle) under the indicated treatment conditions. No significant changes in viability were observed across all groups, indicating that the effects of **LT12** are not attributable to cytotoxicity. Data are presented as mean ± SD (n = 5). Statistical significance was determined by one-way ANOVA with appropriate multiple-comparisons tests. ns denotes not significant.

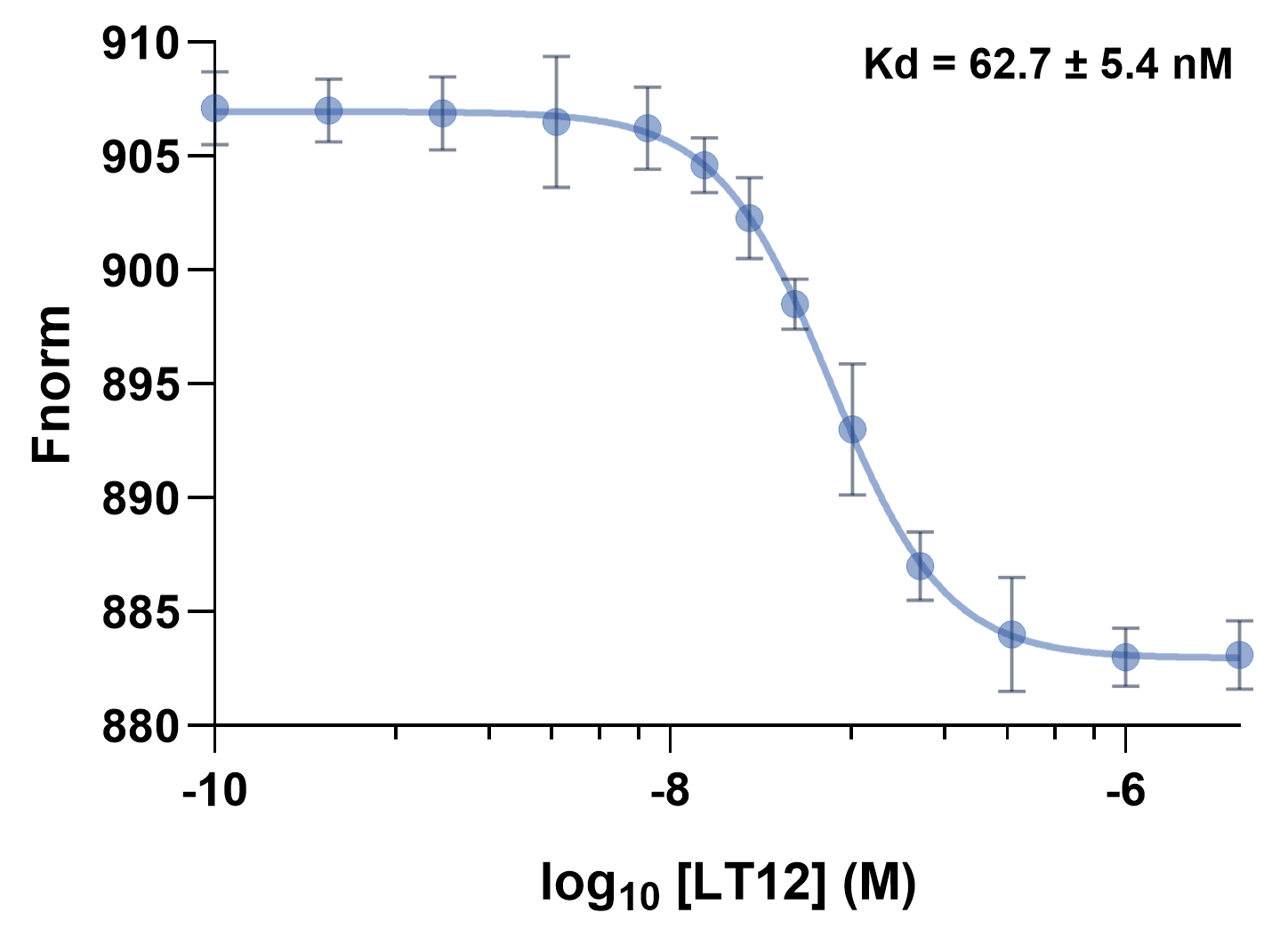

**Figure S9. Cross-species binding of LT12 to murine ILT3 measured by MST.** Normalized thermophoresis signals were recorded across a concentration series of **LT12** against the recombinant extracellular domain of mouse ILT3. Data shows a concentration-dependent change in thermophoretic response consistent with specific binding. Data are presented as mean ± SD (n = 5).

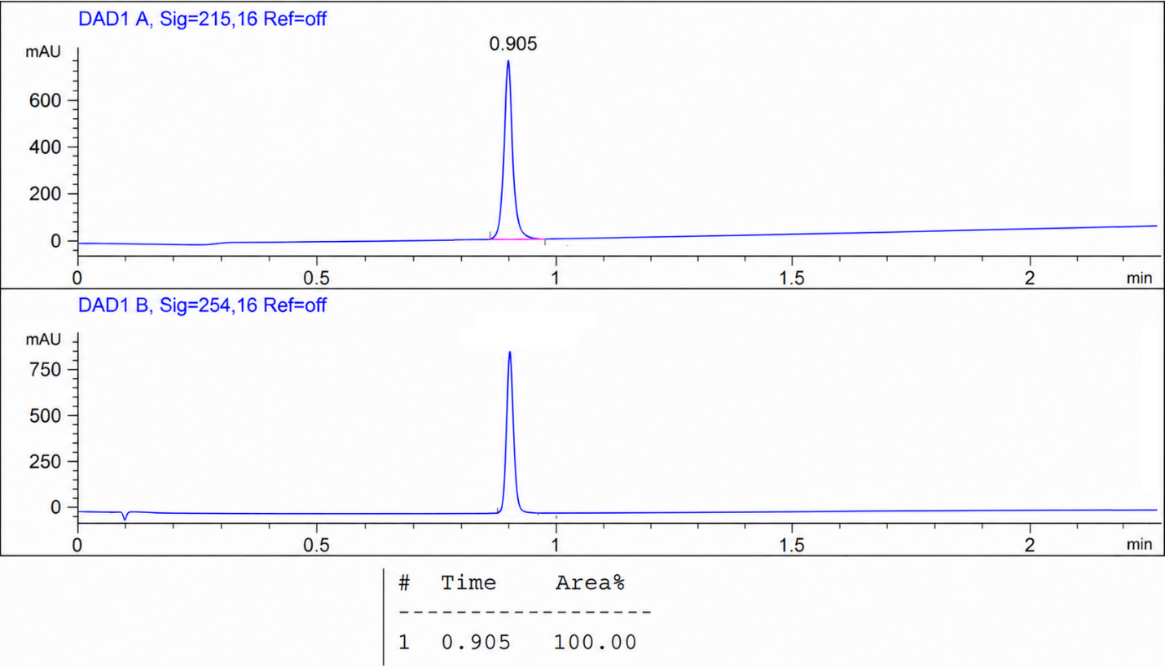

**Figure S10.** HPLC trace for compound **LT12**.
